# The genetic architecture of cell-type-specific cis*-*regulation in maize

**DOI:** 10.1101/2024.08.17.608383

**Authors:** Alexandre P. Marand, Luguang Jiang, Fabio Gomez-Cano, Mark A.A. Minow, Xuan Zhang, John P. Mendieta, Ziliang Luo, Sohyun Bang, Haidong Yan, Cullan Meyer, Luca Schlegel, Frank Johannes, Robert J. Schmitz

## Abstract

Gene expression and complex phenotypes are determined by the activity of cis-regulatory elements. However, an understanding of how extant genetic variants affect cis regulation remains limited. Here, we investigated the consequences of cis-regulatory diversity using single-cell genomics of >0.7 million nuclei across 172 *Zea mays* (maize) inbreds. Our analyses pinpointed cis-regulatory elements distinct to domesticated maize and revealed how historical transposon activity has shaped the cis-regulatory landscape. Leveraging population genetics principles, we fine-mapped ∼22,000 chromatin accessibility-associated genetic variants with widespread cell-type-specific effects. Variants in TEOSINTE BRANCHED1/CYCLOIDEA/PROLIFERATING CELL FACTOR binding sites were the most prevalent determinants of chromatin accessibility. Finally, integrating chromatin accessibility-associated variants, organismal trait variation, and population differentiation revealed how local adaptation has rewired regulatory networks in unique cellular context to alter maize flowering.

## Main Text

Understanding the genetic and molecular principles underlying quantitative trait variation is paramount to improving crop yield and climate resiliency. Towards this goal, genome-wide association studies (GWASs) of genetically distinct inbred *Zea mays* (maize) diversity panels have identified thousands of genetic variants associated with agriculturally important characteristics (*1*). However, due to linkage disequilibrium (LD), disentangling large numbers of linked genetic variants to derive causal effects remains challenging. In maize, trait-associated variants are mostly non-genic, and lie ∼16.4 kb from the nearest gene, complicating efforts to infer the molecular mechanisms contributing to phenotypic diversity (*2*). Thus, complementary approaches are required to pinpoint causal GWAS variants.

Variation in the patterns and magnitude of gene expression is a significant contributor to quantitative phenotypes. Spatiotemporal patterning and kinetics of gene expression is orchestrated by cis-regulatory elements (CREs), short DNA sequence motifs bound by sequence-specific transcription factors (TFs). Investigation of CREs has been useful in furthering the understanding of the molecular determinants behind phenotypic manifestation (*3–5*). Indeed, studies have illustrated that CRE sequence evolution leads to morphological innovation and species divergence (*6, 7*). Although some CRE variants have been associated with phenotypic changes (*8–13*), the mechanistic links between CRE variation and phenotypic diversity remain unclear. Moreover, different cell types are hallmarked by distinct CRE usage that is licensed by differential chromatin accessibility. As a result, how regulatory genetic variants affect molecular processes in a cell-type dependent manner remains unclear. Thus, defining the cellular contexts in which GWAS CRE variants act is a powerful avenue to understand polygenic trait manifestation (*14*).

Compared to humans (r^2^=0.2 at 50-kb) (*15*) and mice (r^2^=0.2 at 200-kb) (*16*), LD in the maize Goodman-Buckler diversity panel (*17*) is substantially lower (r^2^=0.2 at ∼1.4-kb; **fig. S1**), presenting a unique opportunity to uncover the causal variants associated with molecular phenotypes. Taking advantage of this experimental system, we performed single-nuclei chromatin accessibility profiling in a diverse panel of 172 geographically distributed maize inbreds. We established the TF binding sites (TFBSs) contributing to accessible chromatin region (ACR) sequence conservation within maize and identified transposon-derived CREs that shaped the regulatory landscape of domesticated maize. Exploiting the quantitative nature of chromatin accessibility, we identified more than 100,000 single nucleotide variants (SNVs) within ACRs associated with chromatin accessibility variability (chromatin accessibility quantitative trait loci, caQTL) and fine-mapped more than 22,000 SNVs at cell-type-resolution. Comprehensive investigation of this dataset uncovered genetic variants within TF footprints for TEOSINTE BRANCHED1/CYCLOIDEA/PROLIFERATING CELL FACTOR (TCP) as the most significant determinants of chromatin accessibility and putative chromatin interactions. Finally, by integrating cell state-resolved caQTL with GWAS hits, signatures of population differentiation, and TF binding sites, we shed light on the context-dependent regulatory mechanisms that enabled tropical maize to adapt to temperate climates.

### Widespread intraspecific cis-regulatory variation

To establish the impact of genetic variation on cis-regulatory function at single-cell resolution in maize, we profiled chromatin accessibility from ∼1.37 million seedling nuclei across 172 distinct maize inbred lines using single-cell Assay for Transposase Accessible Chromatin sequencing (scATAC-seq) with a pooling, replication (average ∼3 replicates per genotype), and *in silico* genotyping framework (**Fig. 1A-1C**; **fig. S2**-**S3**; **table S1**-**S2**). Following strict thresholding for genotype calls, we retained 682,794 high-quality nuclei (average of 3,970 nuclei per inbred) and identified a total of 108,843 ACRs (**Fig. 1C**; **fig. S3**; **table S3**). Variant calls and chromatin accessibility profiles were consistent with prior genotyping (average genetic correlation=0.91) and among replicated genotypes (average Spearman’s correlation coefficient, SCC=0.92), respectively (**fig. S3**).

**Fig. 1.**
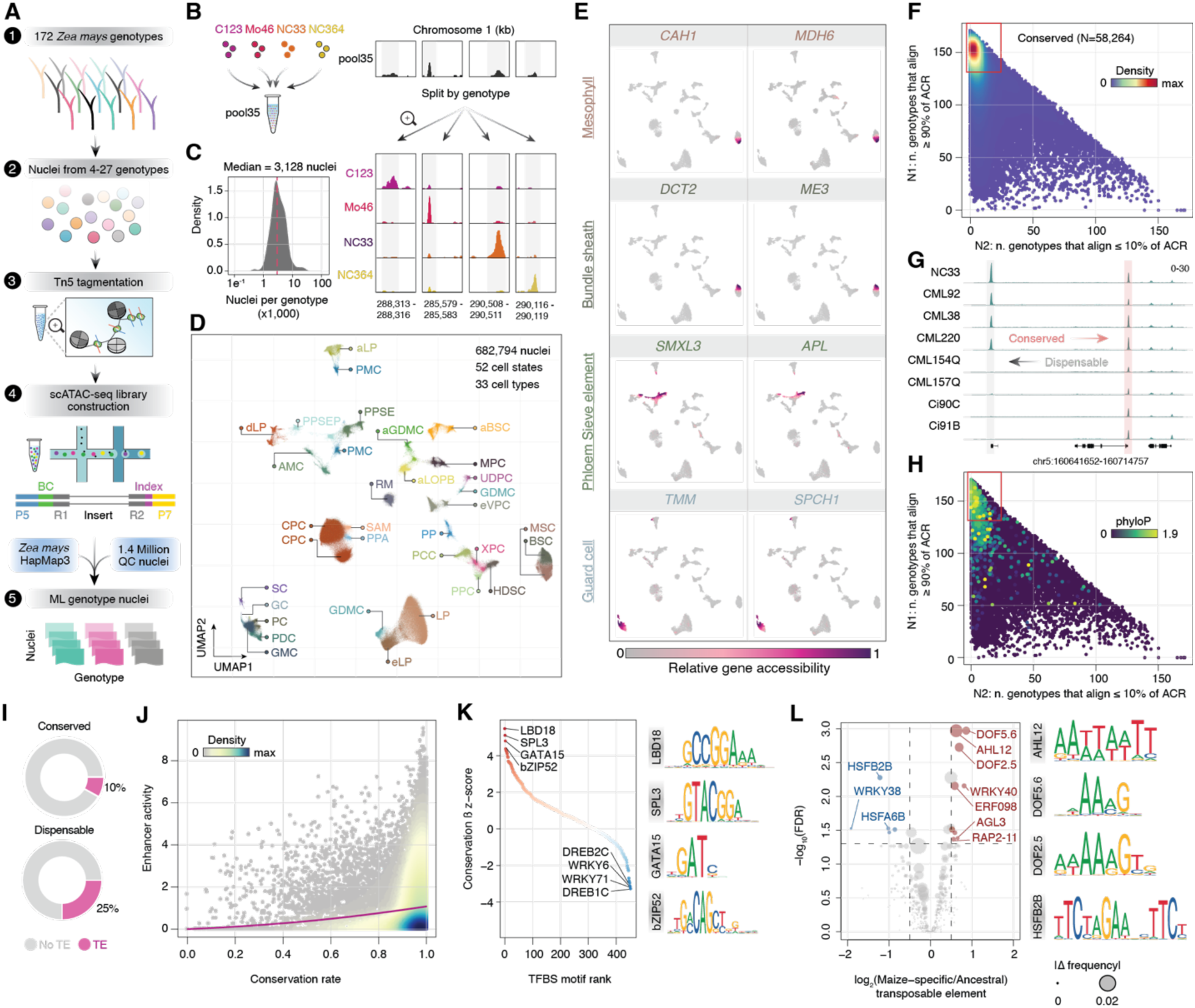
Widespread cis-regulatory sequence variation. (**A**) Experimental and computational workflow. (**B**) Pooling strategy and genotype deconvolution. (**C**) Distribution of nuclei counts per genotype. (**D**) UMAP visualization of chromatin accessibility variance among nuclei from diverse maize genotypes. (**E**) Visualization of chromatin accessibility variance for eight major cell-type-specific marker genes. (**F**) Scatter plot illustrating the number of genotypes with greater than 90% coverage from whole-genome resequencing of an ACR (y-axis) versus the number of genotypes with less than 10% coverage of an ACR (x-axis). Points are colored by density. (**G**) Genome browser examples of conserved and dispensable ACRs identified from panel F. (**H**) Same as panel F, except ACRs are colored by weighted average phyloP scores among grass species. (**I**) Pie charts comparing the proportion of ACRs overlapping transposable elements from the conserved and dispensable classifications. (**J**) Scatter plot of ACR conservation rate (x-axis) and enhancer activity (y-axis) as estimated by STARR-seq. Points are colored by density. (**K**) Ranked scatter plot of permutation normalized TFBS effects towards ACR conservation rate adjacent to exemplary TF motifs. (**L**) Bubble plot of the log_2_ transformation of the ratio between the fraction of maize-specific TE-ACRs and the fraction of ancestrally conserved TE-ACRs containing a given TF motif (x-axis) against the log_10_ transformed FDR. The size of the bubble reflects the absolute difference in TE-ACR frequency for a given TF motif. Exemplary TF motifs are illustrated on the right.

We adopted a reference mapping approach to ensure robust cell-type identification (**fig. S4A**- **S4C**). Specifically, a reference set of high-quality nuclei (n=82,283) from all 172 genotypes were clustered into 52 distinct groups with similar genome-wide chromatin accessibility profiles providing a reference embedding to project the remaining nuclei (**Fig. 1D**). To assign cell identity to each cluster, we estimated gene chromatin accessibility scores for all nuclei and assessed chromatin accessibility dynamics for known marker genes via an iterative strategy composed of (*i*) automated annotation based on cell-type and cell cycle-stage enrichment scores, (*ii*) focused cell-type annotation refinement by evaluating differentially accessible chromatin of known cell type and spatial marker genes, and (*iii*) visual inspection of marker gene enrichment (**Fig. 1E**; **fig. S4D**). The final cell-type, cell cycle-stage, and marker-based spatial labels (hereafter referred to as cell state, n=33) were supported through an integration with 30,305 nuclei generated from single-nucleus RNA sequencing (snRNA-seq) (**fig. S5**), with cell state proportions generally consistent among genotypes (**fig. S6**).

Genome-wide documentation of intraspecific cis-regulatory variation is limited (*18*). To understand the relationship between genetic variation and ACR conservation, we evaluated sequence conservation of 82,098 non-coding ACRs in the context of the B73 reference genome and found 58,264 (71%) ACR sequences conserved among genotypes (**Fig. 1F**-**1G**). The remaining dispensable ACRs were (*i*) associated with significantly lower evolutionary conservation rates (phyloP; 5.74-fold depletion, Wilcoxon rank sum test, WRST; *P*<2.2e^-16^), (*ii*) overrepresented in intergenic and gene-proximal regions, and (*iii*) overlapped significantly with certain repetitive sequences (Fisher’s exact test, FET; *P*<2.2e^-16^; **Fig. 1H**-**1I**; **fig. S7A**-**S7B**; **table S3**). Showcasing the functional importance of conserved ACRs, we found a significant positive relationship between conservation rate and self-transcribing active regulatory region sequencing (STARR-seq) enhancer activity (Likelihood ratio test, LRT; *P*<2.2e^-16^; **Fig. 1J**) (*19*). We hypothesized that selective constraints on the activities of specific TFBSs could underlie ACR conservation. Indeed, a Monte Carlo permutation ridge regression test integrating resequencing information associated the occurrence of 64 TFBS motifs with elevated ACR sequence conservation (FDR<0.05; **Fig. 1K**; **table S4**). Most of these highly conserved TFBS motifs recruit critical developmental regulators, including TFs from *LATERAL ORGAN BOUNDARY DOMAIN* (*LBD*) (*20*), *SQUAMOSA PROMOTER BINDING PROTEIN-LIKE* (*SPL*) (*21*), and *GATA* gene families (*22*). TFBS motifs from the environmentally responsive *WRKY* and *DEHYDRATION RESPONSE ELEMENT BINDING* (*DREB*) gene families exhibited strong negative association with conservation, consistent with stimuli-response CREs contributing to divergence and local adaptation (**Fig. 1K**) (*23, 24*).

Domestication of maize from its wild progenitor, *Z. mays* ssp. *parviglumis* (teosinte), was partially driven by selection of extant cis-regulatory variants (*10, 13, 25*). To identify CREs unique to the transition to cultivated maize, we first isolated ACRs (n=9,174) lacking sequence alignments in 21 teosinte genotypes (**table S3**). Consistent with a recent origin, domesticated maize-specific ACRs were enriched for dispensability (FET, *P<*2.2e^-16^), lower enhancer activity (WRST, *P<*2.2e^-16^), and increased overlap with repetitive elements (FET, *P<*2.2e^-16^; **fig. S7C**- **S7E**). Many domestication-associated regulatory variants were derived from transposable element (TE) sequences (*10, 13, 26, 27*). Therefore, we asked if proliferation of certain TE families could have contributed to domestication by re-wiring TF targets. Of the 9,174 ACRs specific to domesticated maize, 1,587 were fixed across all inbreds. We found a significant enrichment of *hAT* and *PIF/Harbinger* DNA TE families within 483 fixed ACRs post-dating teosinte divergence (Chi-square test, CST; FDR<0.05; **fig. S7F**). The TEs overlapping domesticated maize-specific ACRs contained TFBSs corresponding to DNA-BINDING WITH ONE FINGER (DOF) (*28*), ETHYLENE RESPONSE FACTOR (ERF) (*29*), and SEPALLATA (SEP) (*30*) TFs, which regulate vasculature and floral development (**Fig. 1L**). *PIF/Harbinger* insertions are known to affect the floral transition by modulating vascular expression of *CONSTANS*, *CONSTANS-LIKE*, and *TIMING OF CAB1* (*CCT*) clock genes (*8, 13*), consistent with our genome-wide results. These results highlight that co-option of *hAT* and *PIF/Harbinger* CREs may have played a widespread role in rewiring the regulatory landscape of domesticated maize.

### Genetic determinants of chromatin accessibility

Single-cell data allows investigation of cis-regulatory variation at cell-type resolution. To this end, we performed caQTL mapping by modeling normalized chromatin accessibility as a function of SNV allelic dosage and a suite of latent and measured covariates. We uncovered 4.6 million cis-caQTL associated with 23,959 and 23,123 ACRs (hereafter termed caQTL-ACRs; dynamic beta-adjusted FDR<0.1–0.01) at the bulk-tissue and cell state-resolved scales, respectively (**Fig. 2A**-**2C**; **fig. S8**). As expected, we found that SNVs within ACRs (22%; 107,623/481,947) were enriched for caQTL (FET, odds ratio=1.93, *P*<2.2e-16; **fig. S8F**).

**Fig. 2.**
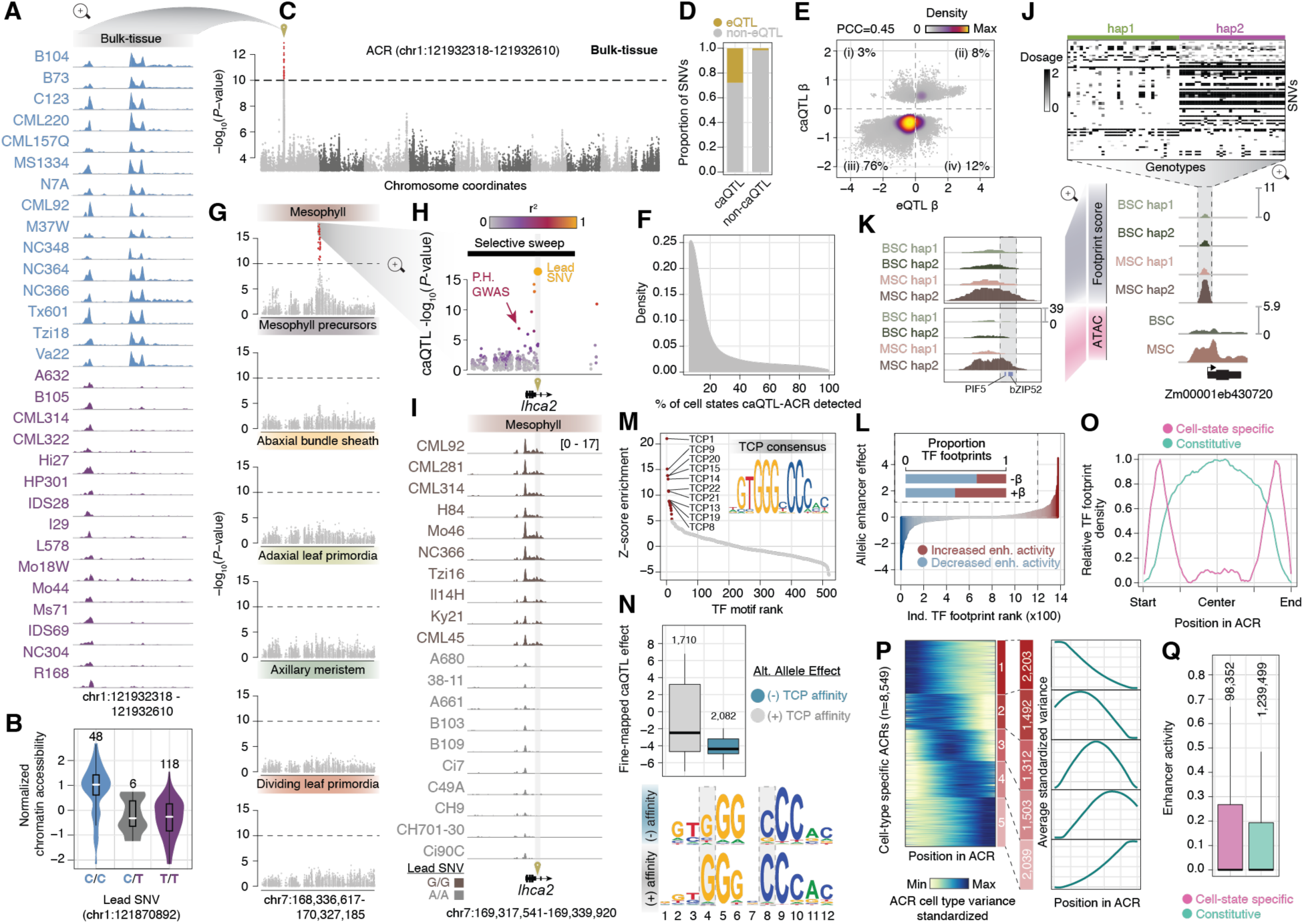
Genetic determinants of chromatin accessibility variation. (**A**) Genome browser view of a caQTL-ACR in 30 exemplary bulk-tissue aggregates colored by the lead SNV genotype in panel B. (**B**) Violin plot of the distribution of normalized chromatin accessibility for three caQTL genotypes associated with the ACR in panel A. (**C**) Manhattan plot of SNV significance by genomic position for a single ACR phenotype in pseudobulk-tissue aggregates. (**D**) Coincidence of caQTL and eQTL between the closest ACR-gene pairs from bulk-tissue association mapping. (**E**) Comparison of caQTL and eQTL effect sizes and directions for SNVs from matched ACR-gene pairs. The percentage of variants are denoted in each quadrant. (**F**) Distribution of caQTL-ACRs detection across cell states. (**G**) Manhattan plot for a cell state-specific caQTL. (**H**) Variant association plot versus genomic position for the ACR shown in panel I. Points are colored by linkage disequilibrium (r^2^) relative to the lead variant. A previous GWAS hit for plant height is shown by the red arrow. The location of a selective sweep is shown as a black bar. (**I**) Genome browser view of 20 exemplary genotypes at the *lhca2* locus aligned with the genomic coordinates of panel H. Tn5 integrations are scaled per million. (**J**) Example of haplotype clustering and TF footprinting in bundle sheath and mesophyll cells. (**K**) Zoom-in on allele-specific footprint scores (top) and chromatin accessibility (bottom). (**L**) Main: Allelic difference in enhancer activity for 1,379 TF footprints. Inset: Proportion of TF footprints with decreased or increased enhancer activity grouped by caQTL effect direction. (**M**) TF footprint motifs ranked by enrichment with fine-mapped caQTL. Red dots indicate FDR < 1e^-5^. (**N**) Top: Distribution minus outliers of fine-mapped caQTL effects for SNVs increasing and decreasing TCP binding affinity. Bottom: Motif logos from variants predicted to increased and decrease TCP binding affinity. (**O**) Distribution of cell state-specific and constitutive TF footprints across ACRs. (**P**) Heatmap illustrating standardized ACR variance across cell types clustered by k-means. The average cluster standardized variances are shown to the right. (**Q**) Distribution minus outliers of enhancer activity for cell state specific and constitutive TF footprints.

Several caQTL-ACRs represented known CRE variants with trait effects, including those regulating *ZmCCT9* (*8*)*, ZmRAP2.7* (*9*)*, gt1* (*12*), and *ZmRAVL1* (*11*) (**fig. S9A**). Additionally, bulk caQTL overlapped significantly (26%; CST, *P<*2.2e^-16^) with previously identified expression QTL (eQTL) (*31*) with ∼85% effect direction concordance (**Fig. 2D**-**2E**; **fig. S9B**-**S9F**). Most alternate alleles (76%) were associated with both reduced chromatin accessibility and transcription, indicating that a majority of alternate allele caQTL disrupt transcription-activating CREs (**Fig. 2E**). However, we also find matched caQTL-eQTL variants associated with increased chromatin accessibility and transcription. These variants were enriched in Polycomb Response Elements (PREs) recognized by the BASIC PENTACYSTEINE (BPC) family of TFs (BPC5, BPC6, and BPC1) relative to background (*32, 33*), suggesting that variants inhibiting BPC-based Polycomb Repressive Complex 2 targeting occur frequently and increase local chromatin accessibility and gene expression (FET, *P*<7.1e^-5^; **fig. S10A**).

We posited that bulk tissue cellular heterogeneity masks caQTL from rare cellular contexts. Indeed, 44% of caQTL-ACRs identified by cell state-resolved caQTL mapping were not found in bulk tissue (**fig. S10B**). These caQTL-ACRs were associated with greater chromatin accessibility specificity (WRST, *P<*3.5e^-112^; **fig. S10C**), suggesting that many caQTL may be cell state-specific. To determine the prevalence of caQTL specificity, we explicitly modeled caQTL across cellular contexts via multivariate adaptive shrinkage (*34*). We found 53% of cell state-resolved caQTL were restricted to three or fewer cell states in contrast to ∼1.2% of caQTL shared across all cellular contexts (**fig. S10D**-**S10E**). Likewise, caQTL identified in select contexts and less abundance cell states were more frequently missed in bulk tissue (**fig. S10F**- **S10G**). Cell state-specific caQTL also exhibited more frequent changes (11%) in effect directions between cell states unlike shared and bulk-tissue caQTL (<1%; **fig. S10H**-**S10J**). Highlighting the utility of single-cell profiling, we found that caQTL-ACRs unique to a cell state often overlapped with variants associated with complex traits (**Fig. 2F**-**2G**). For example, in the last intron of *light harvesting complex a2,* we identified a mesophyll-specific caQTL-ACR missed in bulk tissue in LD (r^2^=0.6) with a GWAS hit for plant height and embedded within a selective sweep in temperate non-stiff stalk germplasm (**Fig. 2H**-**2I**) (*35*). These results highlight that cell state-specific caQTL are common in maize and can underpin trait variance.

### caQTL perturb TFBSs

A total of 107,623 caQTL were physically located within caQTL-ACRs, leading us to hypothesize that causal caQTL perturb TFBSs. Focusing on cell state-resolved caQTL-ACRs, we fine-mapped 22,053 putatively causal caQTL (95% credible sets; average of 3,054 per cell state; **fig S11A**; **table S5**). Next, within each cellular context, we identified differential TF footprints between the two most frequent haplotypes, reflecting alleles with differential TF occupancy (**Fig. 2J**). This analysis revealed an average of ∼25,000 allele-specific TF footprints per cell state, including examples of allele- and cell state-specific chromatin accessibility over TF footprints (**Fig. 2K**; **fig. S11A**). Consistent with functional regulatory elements, TF footprints had higher STARR-seq enhancer activity compared to nearby controls (50-bp shift; WRST; *P<*6.9e^-101^; **fig. S11B**). Nearly half (47%) of fine-mapped caQTL were in TF footprints, a 50-fold enrichment compared to randomized control regions (Monte Carlo permutation, MCP; *P*<1e^-4^; **fig. S11C**). To validate the effects of the fine-mapped variants within TF footprints, we used STARR-seq data to quantify enhancer activity from both reference and alternate alleles.

Fine-mapped TF footprint variants had significantly greater absolute effect sizes on STARR-seq enhancer activity compared to non-caQTL variants (WRST; *P<*2.6e^-73^; **fig. S11D**). Moreover, TF footprint alleles associated with negative effect caQTL had a greater frequency of decreased enhancer activity (CST; *P<*9.8e^-8^; **Fig. 2L**). Cell state-resolved analysis of TF footprint enrichment was consistent with known TF usage among cells states; footprint enrichment was observed for REVEILLE1 (RVE1) in mesophyll (*36*), DNA-BINDING WITH ONE FINGER (DOF) in vasculature (*37*), KANADI (KAN) in abaxial bundle sheath (*38*), and E2F in dividing leaf primordia cells (**fig. S11E**-**S11G**) (*39*). We observed coincidence of cell-state specificity among fine-mapped caQTL, differential TF footprints, and chromatin accessibility, suggesting that variants with cell state-specific effects precisely alter chromatin accessibility by perturbing cell state-specific TF binding (MCP, *P<*1e^-4^; **fig. S11H**). To test this, we assessed the predicted allelic effects of fine-mapped SNVs on TF-binding affinity. Fine-mapped caQTL were associated with significantly greater absolute allele effects compared to non-caQTL SNVs (MCP; *P*<1e^-4^), suggesting that putatively causal caQTL affect chromatin accessibility by influencing TF binding affinity (**fig. S11I**-**S11J**).

We hypothesized that only select TFBS variants affect chromatin accessibility. To this end, we compared the enrichment of TF footprint motifs between fine-mapped caQTL and non-caQTL SNVs. Fine-mapped caQTL within TFBSs recognized by TCP family members had the strongest effects (MCP, FDR<1e^-5^; **Fig. 2M**). Moreover, fine-mapped alleles with reduced TCP binding affinity, particularly at the 4^th^ and 8^th^ base positions, were exclusively associated with strong decreases in chromatin accessibility (WRST; *P<*5.6e^-8^; **Fig. 2N**). Investigating expression patterns of TCP TFs revealed greater cell-state specificity than other TFs, suggesting that distinct TCP TFs promote accessible chromatin in unique cellular contexts (**fig. S12A**-

**S12B**). We reasoned that characterizing regulation of TCP TFs could provide additional context for their role in promoting accessible chromatin. To this end, we generated TF-ACR gene regulatory networks (GRNs) for each cell state using metacells from the integrated scATAC-seq/snRNA-seq embedding (**fig. S12C**). As expected, unique cell states were hallmarked by distinct core TFs based on centrality scores, such as GLOSSY3 in protoderm, GOLDEN2-LIKE TFs in mesophyll, and DOF TFs in procambial cells (**fig. S12D**). While the most connected TFs in each cell state were generally non-TCPs, 90% (18/20) of cell states had at least one TCP TF in the top 10% of TFs ranked by centrality (**fig. S12E**). Moreover, the centrality of these TCPs was highly specific to the cognate cell state (**fig. S12F**). TCPs are regulated by TFs with significantly greater numbers of targets relative to other TFs genes, suggesting cell state-specific TCP transcription is coordinated by master regulators of cell identity (WRST, *P*<2.24e^-35^; **fig. S12G**).

Next, we noted that cell state-specific TF footprints tended to occur on ACR edges (**Fig. 2K**). Indeed, cell state-specific TF footprints were biased towards ACR boundaries compared to TF footprints present across multiple cell states globally (**Fig. 2O**). To measure chromatin accessibility dynamism at ACR boundaries, we estimated and k-means clustered the cell state variance in chromatin accessibility along the length of the ACR sequence for each cell state-specific ACR (n=8,549). Approximately 86% of cell state-specific ACRs were most variable in boundary regions (clusters 1, 2, 4, 5; **Fig. 2P**). Quantifying TF footprint frequencies spatially within ACRs (**fig. S13A**) revealed that the collection of TF footprint motifs in ACR peripheries are highly unique to each cell state (**fig. S13B**). These boundary TFBSs had greater STARR-seq enhancer activity compared to constitutively accessible TFBSs (WRST; *P<*1.2e^-12^; **Fig. 2Q**), indicating dynamic ACR boundaries unveil potent cell state-specific CREs. Consistent across cell states, the top 20 most enriched TF footprints within ACR centers all corresponded to TCP TFs (**fig. S13C**). Taken together, these results suggests that ACRs are licensed or maintained

### Genetic determinants of long-range chromatin interactions

More than 56% of bulk-scale caQTL were associated with more than one ACR (**fig. S14A**), suggesting a single caQTL might impact spatial chromatin interactions. Similar to past studies in humans (*40*), we identified 29,281 long-range caQTL (29,293 ACR-ACR linkages), defined as a caQTL variant associated with chromatin accessibility changes at more than one ACR (**Fig. 3A**). On average, long-range caQTL were associated with 3.1 ACRs and commonly linked gene- distal regions (**fig. S14B**-**S14C**). Relative to randomly permuted caQTL-ACRs links, long-range caQTL were enriched for co-accessible ACRs (MCP, *P*<1e^-4^), predicted chromatin interactions based on a deep learning convolution neural network (MCP, *P*<1e^-4^) (*41*), and HiC and HiChIP chromatin loops (MCP, *P*<1e^-4^; **Fig. 3B**-**3C**; **fig. S14D**). For example, we identified a caQTL within the promoter of *abs*cis*ic acid insensitive 8* (*abi8*), an ABI-VP1 TF with a hypothesized negative regulatory role in leaf blade morphogenesis (*42*). The *abi8* promoter variant was associated with ACRs 10-kb and 50-kb downstream coincident with H3K4me3- and H3K27me3- HiChIP chromatin loops (**Fig. 3D**). Additionally, long-range caQTL with fine-mapped variants showed stronger overlap with chromatin loops than those lacking fine-mapped caQTL (FET; *P*<1.49e^-6^). These long-range fine-mapped caQTL exhibited larger effect sizes and smaller standard errors than non-fine-mapped caQTL, indicating that large-effect caQTL are better powered to detect distal chromatin interactions (**fig. S14E**). As a result, the number of long- range QTL in this study is likely underestimated.

**Fig. 3:**
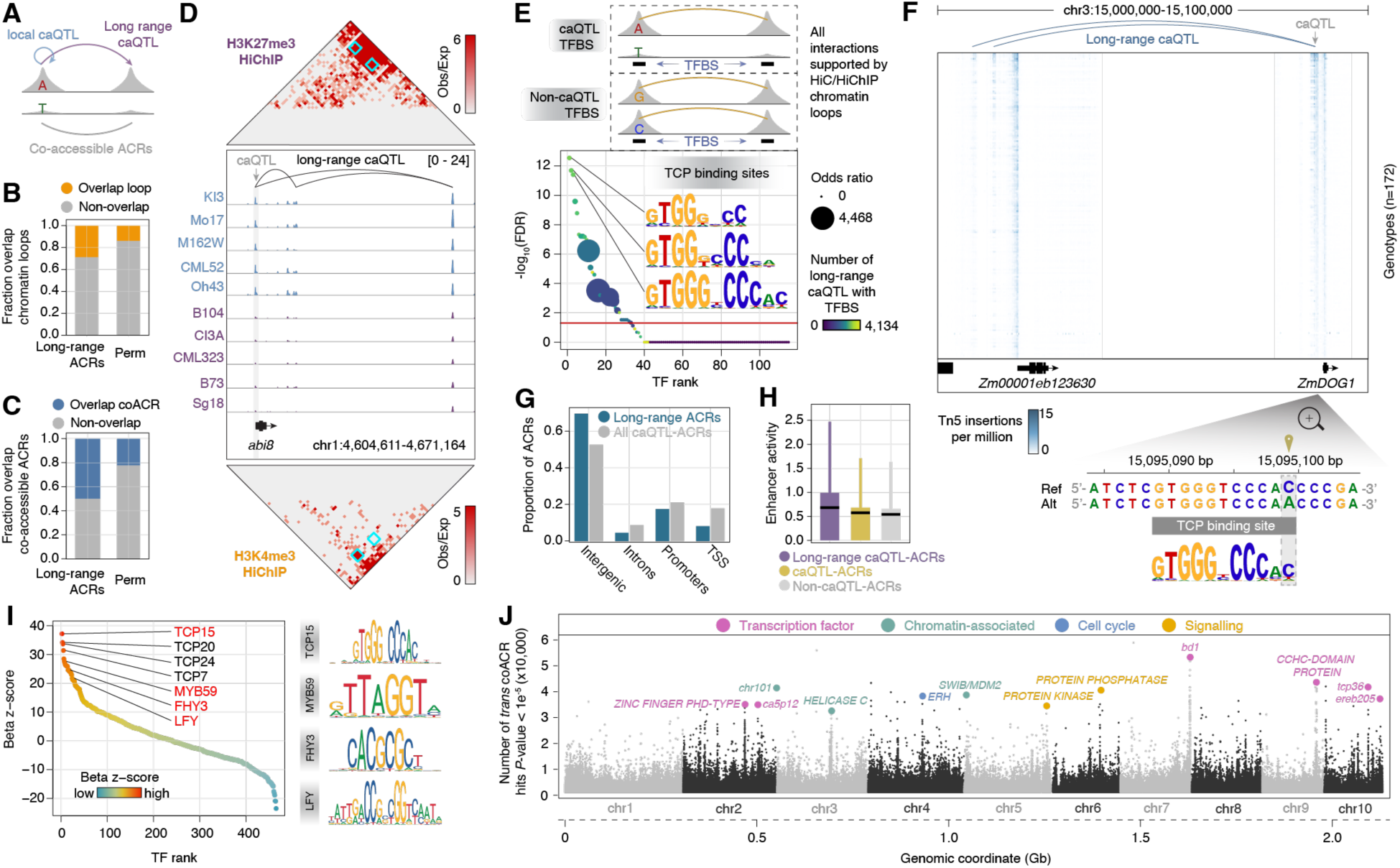
Long-range caQTL perturb chromatin loops. (**A**) Schematic of local and long-range caQTL (top), and co- accessible ACRs (bottom). (**B**) Fraction of predicted (based on long-range caQTL genetic evidence) and permuted ACR-ACR interactions overlapping experimentally resolved chromatin loops. (**C**) Fraction of predicted and permuted ACR-ACR interactions overlapping with co-accessible ACRs. (**D**) Middle: Exemplary long-range caQTL interaction between the promoter of *abi8* and two ACRs downstream. Top: Normalized H3K27me3 HiChIP contact matrix. Bottom: Normalized H3K4me3 HiChIP contact matrix. Blue rectangles denote the interaction frequency between the promoter of *abi8* and the downstream ACRs. (**E**) Top, illustration of test sets (TFBS counts containing long-range caQTL or non- caQTL SNVs conditioned on HiC/HiChIP looping and reciprocal presence of TFBSs at both anchors). Bottom, bubble plot of TF enrichment ranks by -log_10_(FDR). Points are colored by the number of interactions and sized by the odds ratio from Fisher’s exact tests. Red line indicates -log_10_(FDR = 0.05). Inset, TFBS logo of top three enriched motifs corresponding to 17 TCP TF family members. (**F**) Top, density heatmap of Tn5 insertion counts across genotypes (rows). Bottom, zoom-up of caQTL alleles overlapping a predicted TCP binding site. (**G**) Proportion of ACR in each genomic context split by long-range caQTL-ACRs with local caQTL and all caQTL-ACRs. (**H**) Distribution of enhancer activity across long-range caQTL-ACRs, all caQTL-ACRs, and non-caQTL-ACRs. Black bars indicate distribution means. (**I**) TF motifs ranked by beta coefficient Z-scores (relative to 100 permutations) from a generalized linear model of long-range caQTL-ACR interactive capacity. (**J**) Manhattan plot of the number of co-accessible ACRs associated (*P-* value < 1e^-5^) with a single SNV. Candidate genes are annotated by different colors. by TCPs, while dynamic accessible chromatin boundaries are influenced by other cell state- specific TFs.

We then aimed to understand the regulatory basis of long-range caQTL. To do so, we compared TFBSs found in ACRs of long-range interactions supported by chromatin loops that colocalized with either (*i*) fine-mapped caQTL or (*ii*) SNVs with non-significant effects on chromatin accessibility (**Fig. 3E**). Like variants that impact local chromatin accessibility, caQTL affecting long-range interactions were enriched for TCP family TFBSs (**Fig. 3E**). For example, a local caQTL affecting a predicted TCP binding site in the promoter of the maize homolog of *DELAY OF GERMINATION 1* (*DOG1*) was associated with accessible chromatin variation of two distal co-accessible ACRs 88-kb and 90-kb upstream (**Fig. 3F**). Investigating long-range caQTL further, we find that long-range caQTL within intergenic ACRs were more prevalent than other genomic contexts (CST; *P*<1.7e^-221^), associated with a greater number of interactions (interactive capacity), and had stronger STARR-seq enhancer activity than ACRs near genes (WRST; *P<*2.0e^-28^; **Fig. 3G**-**3H**; **fig. S14F**). To evaluate whether specific TFBSs affect interactive capacity, we fit a generalized linear model of interactive capacity as a function of motif occurrences within long-range caQTL-ACRs and compared the model coefficients with 100 random permutations. TFBSs from the TCP family were strongly associated with increased interactive capacity (**Fig. 3I**). These results implicate gene-distal enhancers containing TCP binding sites as potential determinants of long-range chromatin looping.

Identifying the *trans*-regulators driving chromatin interactions remains challenging in plants. As an orthogonal approach, we reasoned that co-accessibility scores between distal ACRs could be used as a proxy for chromatin interactions. Using co-accessibility scores as a response variable, we performed QTL mapping for co-accessible ACRs supported by HiC/HiChIP loops (n=802,504). This analysis revealed 12 loci associated with genome-wide co-accessibility (**Fig. 3J**), including six transcription factors, three chromatin-associated proteins, two phospho-signaling proteins, and a cell-cycle associated gene (**table S6**). We identified a haplotype containing *tcp-transcription factor 36* (*tcp36*) on chromosome 10 as one of the top candidates associated with variation in genome-wide ACR co-accessibility (**fig. S14G**). The coding sequence for *tcp36* contained several missense mutations and a premature stop codon (**fig. S14H**). These results provide additional support for the association between TCP TFs and global chromatin accessibility variation.

### Selection on regulatory variants shaped complex traits

To understand the relative contribution of regulatory variation towards organismal phenotypes and define the regulatory modules underlying complex traits, we performed chromatin accessibility genome-wide association (CAWA) and transcriptome-wide association mapping (TWA). As expected, phenotypes clustered by physiological relationships and trait associations captured known gene-trait interactions (**Fig. 4A**). For example, transcript abundances and regulatory region chromatin accessibilities for *LONELY GUY 7* (*ZmLOG7*)*, CUP-SHAPED COTYLEDON 2* (*ZmCUC2*), and *knotted 1* (*kn1*) were associated with shoot apical meristem (SAM) morphology (*43*), leaf width (*44*), and ear, tassel, and leaf architecture (*45*), respectively.

**Fig. 4:**
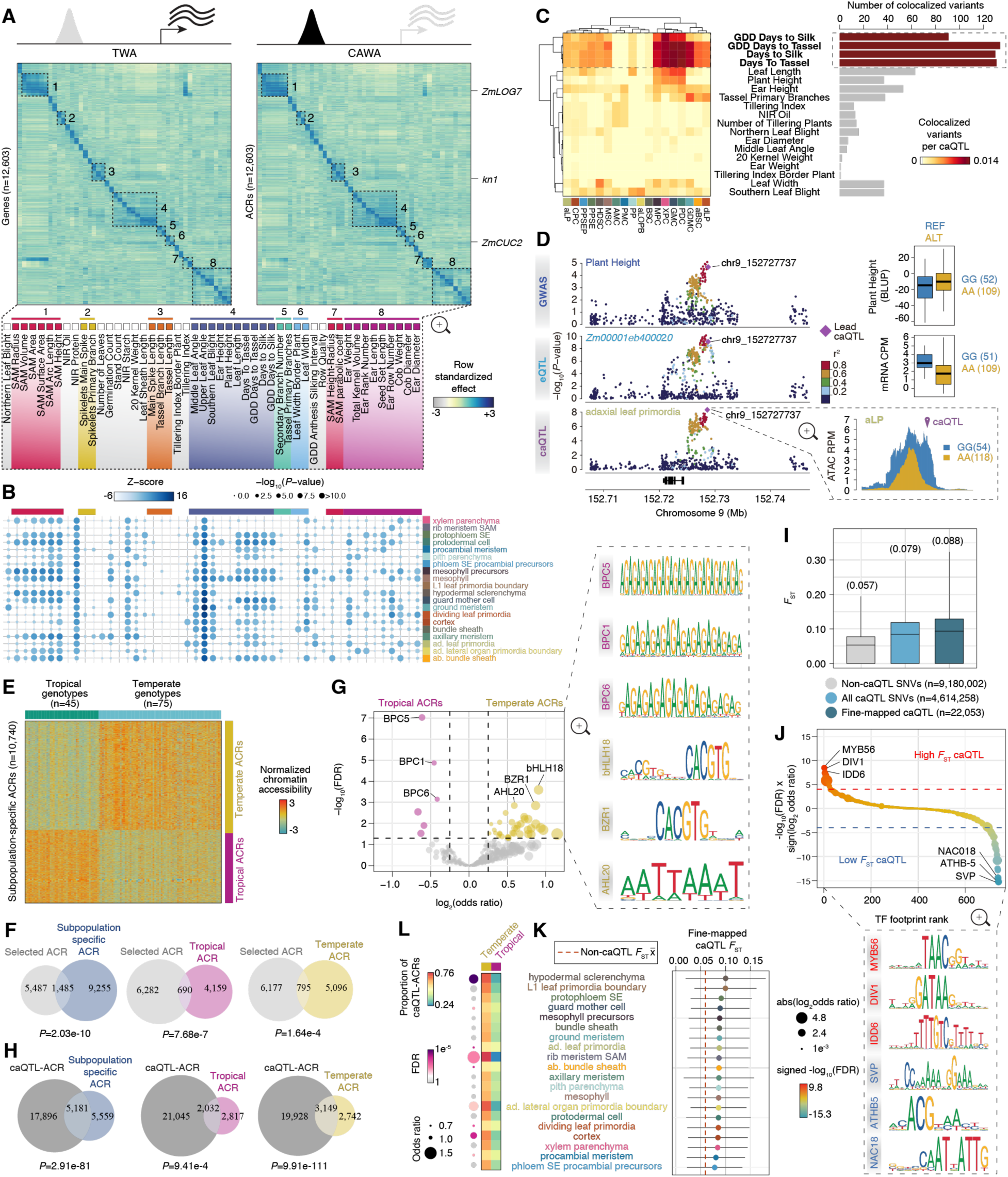
Integration of caQTL with eQTL and GWAS of complex traits. (**A**) Heatmaps of standardized effects from CAWAS and TWAS for matched ACRs (left) and genes (right). Phenotype traits of the columns (identical order for both heatmaps) are shown below. (**B**) Permutation tests (x10,000) of percent variance explained by caQTL partitioned by cell state compared to non-caQTL SNVs. (**C**) Left; heatmap illustrating the fraction of caQTL colocalizing with a given GWAS trait. Right; bar plot of the number of colocalized variants for each GWAS trait. (**D**) Manhattan plots of plant height GWAS (top), expression QTL of bulk seedlings (middle), and caQTL for adaxial leaf primordia (bottom). Variants are colored by LD to the lead caQTL. Trait distributions split by the lead variant genotypes are shown to the right. (**E**) Heatmap of subpopulation-specific normalized chromatin accessibility (rows) across 120 tropical and temperate maize genotypes. (**F**) Venn diagram of ACRs within genomic regions under selection that overlap subpopulation-specific ACRs, tropical-specific ACRs, and temperate-specific ACRs. (**G**) Volcano plot of differentially accessible motifs between tropical and temperate ACRs. (**H**) Venn diagram of caQTL-ACRs that overlap subpopulation-specific ACRs, tropical ACRs, and temperate ACRs. (**I**) Distribution of *F*_ST_ values of non-caQTL SNVs, all caQTL SNVs, and fine-mapped caQTL SNVs. (**J**) TF footprint ranked by the signed significance (-log_10_ FDR) for enrichment of high or low *F*_ST_ SNVs. (**K**) The distribution (standard deviation) and mean *F*_ST_ values of fine-mapped caQTL split by cell state. The dashed red line indicates the genome-wide average for non-caQTL SNVs. (**L**) Heatmap illustrating the proportion of caQTL-ACRs from each cell state that are temperate or tropical specific. Cell states with significant deviations from expected proportions (FDR < 0.05) are denoted by colored bubbles, where the size of the bubble indicates the odds ratio.

CAWA deconvolution identified the TFBSs with the largest impact on organismal traits; SPL and homeodomain motifs associated with SAM and ear morphology, WRKY motifs with flowering time traits, and AUXIN RESPONSE FACTOR (ARF) motifs with leaf width (**fig. S15A**).

Considering complex traits are the synergistic outcome of cellular phenotypes, we compared the percent of trait variation explained by cell state-resolved fine-mapped caQTL relative to permutations of non-caQTL SNVs. Fine-mapped caQTL explained a significant proportion of trait variance for 96% (47/49) of complex traits (**Fig. 4B**). Moreover, organismal traits were often influenced by multiple cell states at varying degrees, indicating cell-distinct processes impact phenotypic variation non-uniformly (**Fig. 4B**). Highlighting the utility of partitioning signal by cell state, caQTL identified only by cell state-resolved profiling explained a greater proportion of trait variation for 29% (14/49) of complex traits relative to bulk caQTL (**fig. S15A**). Furthermore, fine-mapped caQTL from meristem and primordia cells explained the most variance for 69% (34/49) of traits (**fig. S15B**-**S15C**), including flowering time and SAM, ear, and tassel morphologies, supporting prior observations that adult architectural traits associate with regulators of SAM identity (*46*).

To dissect the confluence of chromatin and trait genetic architectures, we used statistical colocalization to identify shared causal variants, revealing 811 caQTL-GWAS trait pairs (**fig. S16**; **table S7**). The most abundant colocalized caQTL-GWAS variants were related to flowering traits (**Fig. 4C**). For example, the alternate allele of a SNV (chr1:261397272) located inside an ACR 5-kb upstream of *lbd10* was associated with decreased chromatin accessibility in epidermal cells and an increase of ∼30 growing degree days to tassel (**fig. S15D**). However, as many causal variants are likely not represented in our study (such as unseen insertions or deletions), we reasoned that finding shared molecular QTL in strong LD (r^2^ > 0.8) with trait-associated variants could be useful for investigating trait regulatory mechanisms. To this end, we uncovered an additional 326 unique caQTL-eQTL variants in LD with 1,342 GWAS hits across 20 cellular contexts (**table S7**). Highlighting the utility of this method to resolve regulatory mechanisms, we identified a GWAS hit for plant height in LD (r^2^=0.93) with a shared adaxial leaf primordia caQTL and eQTL ∼35-kb upstream of the heat shock gene, *hsp19* (Zm00001eb400020; **Fig 4D**). The reference allele was associated with agronomically-favorable decreased plant height, greater *hsp19* transcript abundance, and an expanded accessible chromatin domain. Investigating integrated molecular QTL-GWAS variants more generally, we found that shared caQTL-eQTL were most frequently in LD with GWAS hits for flowering-related traits (**fig. S15E**). Thus, we hypothesized that flowering variability in the diversity panel may be due to regulatory adaptation.

Many physiological processes, including flowering, changed during the migration of maize from tropical to temperate latitudes. Comparing tropical and temperate maize lines, we found 10,740 ACRs with differential accessibility and refer to these regions as subpopulation-specific ACRs (FDR<0.1; **Fig. 4E**). Subpopulation-specific ACRs were strongly enriched in signatures of selection that accompanied the adaptation to temperate climates and GWAS variants associated with flowering time (**Fig. 4F**; **fig. S15G**; **table S8**) (*47*). Moreover, different TFBSs were enriched in subpopulation-specific ACRs depending on the subpopulation. For example, we found transcription-silencing motifs recognized by BPC TFs (*32*) were uniquely associated with tropical ACRs (**Fig. 4G**; **table S9**). In contrast, BRASSINAZOLE-RESISTANT 1 (BZR1), a master developmental regulator, AT-HOOK MOTIF CONTAINING NUCLEAR LOCALIZED 20 (AHL20), a DNA-binding protein associated with delayed flowering (*48*), and a BASIC HELIX-LOOP-HELIX 18 (bHLH18) TF specifically expressed in floral tissue (*49*) were distinctly associated with temperate ACRs (**Fig. 4G**; **table S9**). These TFBS enrichments are consistent with a modified floral transition being integral for temperate adaptation. Despite including genetic relatedness covariates in caQTL mapping, subpopulation-specific ACRs were strongly coincident with caQTL-ACRs (**Fig. 4H**), suggesting that temperate/tropical adaptation underlies a significant proportion of chromatin accessibility variation. Indeed, allele frequency comparisons between tropical and temperate inbreds revealed higher *F*_ST_ values for caQTL and fine-mapped caQTL compared to non-caQTL SNVs (WRST, *P<*2.2e^-16^; **Fig. 4I**). These results highlight prevalent stratifying selection on CRE functional diversity within maize subpopulations.

To define the regulatory grammar promoting temperate and tropical differentiation, we compared TF footprints localized with fine-mapped caQTL categorized as high or low *F*_ST_. High *F*_ST_ fine-mapped caQTL were enriched for TF footprints previously associated with flowering time (MYB56, IDD6) (*50, 51*) and floral cell identity (DIV1) (*52*), and depleted of TF footprints associated with short vegetative phase (SVP) (*53*), ABA-responsiveness (ATHB-5) (*54*) and embryogenesis (NAC018) (*55*) (**Fig. 4J**; **table S10**). We hypothesized that local adaptation would impact chromatin accessibility in specific cell states. Indeed, *F*_ST_ distributions for fine-mapped caQTL differed among cell states and were heavily biased towards ACRs with increased temperate chromatin accessibility, particularly in the rib meristem and hypodermal sclerenchyma (Kruskal-Wallis Rank Sum Test, KWRST, *P<*9.4e^-12^; **Fig. 4K**-**4L**). These results suggest that temperate adaptation involved widespread changes to chromatin accessibility that rewired flowering and developmental TF networks non-uniformly across cell states.

## Discussion

Variable chromatin accessibility underlies cell-type-specific transcriptional regulation that drives functionally diverse cellular states. Although we previously established widespread chromatin accessibility variation among cell states within maize (*56*), the extent to which cell-type level chromatin accessibility differed between genetically diverse individuals remained unclear. Here, we present a large-scale investigation of chromatin accessibility variation in 20 cell states across 172 geographically dispersed maize inbreds. Exploiting low LD in maize, we fine mapped >22,000 putatively causal variants. Fine-mapped caQTL are highly cell state-specific and strongly biased in TCP TFBSs. The patterns of TCP gene expression, association with global co-accessibility, and significant effects on local chromatin accessibility collectively implicates TCP TFs as regulators of chromatin accessibility. Identification of TCP TFs as prevalent and potent promoters of chromatin accessibility and contributors towards chromatin interactions adds to a growing body of evidence implicating TCP TFs as major regulators of chromatin architecture (*57*). Our GRN analysis also found cell state-specific TCPs are targeted by master cell identity TF regulators, showcasing the upstream factors that are important for promoting chromatin accessibility in diverse cell contexts. Identification of TCPs as promoters of accessible chromatin is a potentially exciting advance; only a handful of plant pioneer factors have been identified whilst all studied plants lack a homolog to the metazoan-specific chromatin looping factor, CTCF. We anticipate follow-up genetic and biochemical experiments to investigate the mechanistic roles of TCP factors in chromatin accessibility and long-distance chromosome looping.

Integrating caQTL with GWAS hits uncovered the cellular contexts associated with phenotypic traits. Further, the combination of CAWA, TWA and GWAS provided preliminary insights into the underlying regulatory mechanisms behind complex trait variation. Examining the ACR differences between temperate and tropical maize, we demonstrated that caQTL are associated with signatures of population differentiation and selection, confirming regulatory variation as a key target of local adaptation and human breeding efforts. As such, this resource provides exciting avenues for follow-up exploration and direct real-world application. First, by identifying over 22,000 cis-variants predicted to causally affect chromatin accessibility – of which ∼1,300 are in LD with quantitative phenotypes – we narrow candidate molecular breeding and genetic engineering targets for crop improvement. Second, we revealed how individual TFBS variants affect both the cell-state specificity and enhancer activity of TFBSs, which can inform synthetic CRE design. Third, we defined the ACRs and cognate CREs that are fixed in domesticated maize. The absence of extant genetic variation over these regions both highlights their agronomic importance and occludes their study through traditional quantitative genetic techniques. As such, these fixed regions represent potent exploratory targets for engineering allelic variation via gene editing, both to understand their function and to shepherd novel crop phenotypes. Future reverse genetic and biochemical exploitation of these data will undoubtedly shape crop improvement and deepen mechanistic understanding of phenotypic variation.

Our study addressed several outstanding questions, serving as a foundation for similar caQTL studies. However, we profiled caQTL in only 20 of 33 cell states due to limited power in rare cell types. With technological increases in throughput, future studies will better resolve the caQTL from low-frequency cell types. Additionally, we sampled 7-day old seedlings; this tissue was selected because it includes the shoot apical meristem which is known to forecast adult traits.

However, our sampling strategy cannot find caQTL specific to other phenotypically relevant organs including roots, inflorescences, and seeds, which is also complicated by the small effect sizes of complex traits mapped in our study. These challenges ultimately impact the recovery of colocalized caQTL with known GWAS hits. Expanding the number of tissues and developmental stages profiled by scATAC-seq and snRNA-seq, as well as expanding the number of individuals, will be useful to more comprehensively characterize the molecular underpinnings of trait variance. Another consideration is the use of a single reference genome; our analyses are limited to ACRs and caQTL that can be identified in the context of B73. We anticipate that advances in pangenomics and additional reference genomes will facilitate more comprehensive future discovery (*58*). Finally, although the size of the diversity panel was well-powered to identify cis-caQTL, it was limited for *trans-*QTL detection; uncovering *trans*-regulatory interactions remains a challenge in most eukaryotic genomes. Future studies with larger cohorts and datasets that pre-identify *trans-*regulated ACRs provide a promising avenue to reveal hidden *trans-*QTL.

## Materials and methods summary

Replicated material was collected from ca. seven day old seedlings from 172 diverse maize inbreds from the Goodman-Buckler maize diversity panel (*17*) across several randomized batches (**table S2**). Nuclei were isolated using a previously described approach via flow cytometry sorting (*59*). Flow-sorted nuclei were spun in a swinging bucket centrifuge (5 minutes, 500 rcf) and resuspended in 10 uL of LB01, visualized and counted on a hemocytometer with a fluorescence microscope, and adjusted to a final concentration between 3,200-12,000 nuclei per uL using diluted nuclei buffer (DNP; 10X Genomics, catalog: 1000176). For each library preparation, 5 uL of suspended nuclei were loaded per well on the Next GEM Chip H (10X Genomics, catalog: 1000162). Single-cell ATAC-seq libraries were prepared according to the manufacturer’s instructions (10X Genomics, catalog: 1000176, Chromium Next GEM v1.1) using the Chromium Controller (10X Genomics, catalog: 120223). The resulting scATAC-seq libraries were sequenced using an Illumina S4 flow cell (NovaSeq 6000) in dual-index mode. Raw sequencing output was processed with Cell Ranger Single Cell Software (v2.2.0; 10x Genomics) and aligned to the maize AGPv5 reference genome (*60*) with *chromap* (v0.2.3) (*61*).

Singlet nuclei were identified by removing barcodes associated with multiple genotype identities from *souporcell,* a minimum of 500 unique Tn5 insertions within ACRs, and fraction of reads in peaks and fraction of reads near transcription start sites above background levels determined for each library independently. Cell state identities were identified by analysis of marker genes, differential gene chromatin accessibility, and integration with tissue-matched snRNA-seq data. We then generated chromatin accessibility profiles in each cell state by aggregating nuclei by genotype and cell state identity. Sample covariates were estimated based on genetic relatedness, library pooling information, cell cycle stage, and latent characteristic present in the chromatin accessibility counts matrix. Chromatin accessibility QTL were detected with *tensorQTL* (v1.0.9) (*62*) for each distinct cell state and in the pseudobulk data sets using cell context-specific FDR thresholds based on average read counts.

## Acknowledgements

We wish to thank the GACRC for providing valuable computational support and resources, USDA GRIN for maize germplasm, the Duke University School of Medicine Sequencing and Genomic Technologies Shared Resource for Illumina sequencing services, Dr. Ed Buckler for advice on genotyping strategies, Dr. Joseph Gage for modeling approaches, and Dr. Nathan Springer for feedback on analyses.

## Author contributions

Conceptualization: A.P.M., R.J.S.

Methodology: A.P.M., X.Z.

Investigation: A.P.M., L.J., F.C.G., M.A.A.M., J.P.M., Z.L., S.B., C.M., L.S., F.J., R.J.S.

Visualization: A.P.M., L.J., F.C.G.

Funding acquisition: A.P.M., R.J.S.

Project administration: A.P.M., R.J.S.

Supervision: A.P.M., R.J.S., F.J.

Writing – original draft: A.P.M., M.A.A.M., R.J.S.

## Data and materials availability

All data supporting the conclusions of this study have been made available within the article and the supplementary information. Raw and processed data generated by this study are available in the National Center for Biotechnology Information Gene Omnibus (NCBI GEO) under accession number GSE275410. Coverage tracks from the different inbred lines can be viewed on JBrowse (https://epigenome.genetics.uga.edu/PlantEpigenome/?data=maize_v5). Scripts and software used in the analysis of the data generated by this study are publicly available at https://github.com/plantformatics/maize_282_diversity_scATAC and Zenodo (*63*). Additional processed data sets are available on Dryad (https://doi.org/10.5061/dryad.nk98sf82v).

## Supplementary Materials

### Materials and Methods

#### Plant material and growth conditions

Kernels for 290 maize genotypes were acquired from USDA ARS-GRIN (https://npgsweb.ars-grin.gov). Five biological replicates were sown in 1-inch plug trays with Sungro soil (Sungro Horticulture Canada Ltd) for each genotype in a walk-in growth chamber. Seedlings were maintained under 16-hour 50/50 mixture of 4100K (Sylvania Supersaver Cool White Delux F34CWX/SS, 34W) and 3000K (GE Ecolux w/ starcoat, F40CX30ECO, 40W) fluorescent lighting 50 cm above soil for 7 days at approximately 25°C during light hours with a relative humidity of approximately 54%. Of the 290 genotypes, 177 had germinated after 7 days, the remaining genotypes were excluded from the study. A single representative seedling was selected for each genotype per pool and partitioned to 8 mm either side of the first visible node (16 mm total) with a #2 razor. The 16 mm leaf roll section were rinsed with deionized water and pooled according to the experimental design in **table S2**. All replicates are biological. Samples were collected between 8 and 9 AM (∼1-2 hours after the initiation of the light cycle).

#### Single-cell ATAC-seq library construction

For each multiplexed pool, leaf roll sections were suspended in 500 uL of LB01 buffer (15mM Tris pH 7.5, 2mM EDTA, 0.5mM Spermine, 80mM KCl, 20mM NaCl, 15mM 2-mercaptoethanol, 0.15% TrixtonX-100) and homogenized with a No. 2 straight-edge razor for 120 seconds. The lysate was filtered through two layers of miracloth (Millipore #475855), stained with DAPI to a final concentration of ∼ 1uM and loaded onto a Beckman Coulter Moflo XDP flow cytometer instrument. Between 60,000-120,000 nuclei were sorted from each pool into 1.5mL Eppendorf LoBind® centrifuge tube containing ∼175-350 uL of LB01. Sorted nuclei were spun in a swinging bucket centrifuge (5 minutes, 500 rcf) and resuspended in 10 uL of LB01, visualized and counted on a hemocytometer with a fluorescence microscope, and adjusted to a final concentration between 3,200-12,000 nuclei per uL using diluted nuclei buffer (DNP; 10X Genomics, catalog: 1000176). For each pool, 5 uL of suspended nuclei were loaded per well on the Next GEM Chip H (10X Genomics, catalog: 1000162). Single-cell ATAC-seq libraries were prepared according to the manufacturer’s instructions (10X Genomics, catalog: 1000176, Chromium Next GEM v1.1) using the Chromium Controller (10X Genomics, catalog: 120223). scATAC-seq libraries were sequenced using an Illumina S4 flow cell (NovaSeq 6000) in dual-index mode with eight and 16 cycles for i7 and i5, respectively.

#### Single-nucleus RNA-seq library construction

The protocol for nuclei isolation and purification was adapted from the previously described scifi- ATAC-seq method (*64*). In summary, to minimize RNA degradation and leakage, the same tissue used for scATAC-seq from two maize genotypes, B73 and Mo17, was finely chopped on ice for approximately 1 minute using 600 μL of pre-chilled Nuclei Isolation Buffer containing 0.4U/μL RNase inhibitor (Roche, Protector RNase Inhibitor, Cat. RNAINH-RO) and a comparatively low detergent concentration of 0.1% NP-40. Following chopping, the entire mixture was passed through a 40-μm cell strainer and then subjected to centrifugation at 500 rcf for 10 minutes at 4°C. The supernatant was carefully decanted, and the pellet was resuspended in 500 μL of NIB wash buffer, comprising 10 mM MES-KOH at pH 5.4, 10 mM NaCl, 250 mM sucrose, 0.5% BSA, and 0.2U/μL RNase inhibitor. The sample was filtered again, this time through a 20-μm cell strainer, and gently layered onto the surface of 1 mL of a 35% Percoll buffer. The Percoll buffer was prepared by mixing 35% Percoll with 65% NIB wash buffer in a 1.5-mL centrifuge tube. The nuclei were then subjected to centrifugation at 500 rcf for 10 minutes at 4°C. After centrifugation, the supernatant was carefully removed, and the pellets were washed twice in 650 μL of Nuclei Wash & Resuspension buffer (NWR buffer: 1% BSA, 0.2U/μL RNase inhibitor, 1xPBS). The nuclei were resuspended in 50 μL NWR buffer.

Approximately 5 μL of nuclei were diluted tenfold and stained with DAPI (Sigma Cat. D9542). Subsequently, nuclei quality and density were evaluated with a hemocytometer under a fluorescence microscope. The original nuclei were further diluted with DNB buffer to achieve a final concentration of 1,000 nuclei/μL. Ultimately, a total of 16,000 nuclei were used as input for snRNA-seq library preparation.

For scRNA-seq library preparation, we employed the Chromium Next GEM Single Cell 3’GEM Kit v3.1 from 10X Genomics (Cat# PN-1000123), following the manufacturer’s instructions (10xGenomics, CG000315_ChromiumNextGEMSingleCell3-_GeneExpression_v3.1_DualIndex_RevB). The libraries were subsequently sequenced using the Illumina NovaSeq 6000 in dual-index mode with 13 cycles for the i7 and i5 indices, respectively.

#### scATAC-seq raw data processing

scATAC-seq binary base call sequences files (BCL) output from the Illumina S4 NovaSeq 6000 were demultiplexed with *cellranger-atac mkfastq* (v1.2) and aligned to the B73 AGP v5 maize reference genome (*60*) using *chromap* (v0.2.3) with non-default settings (-l 2000 -q 1 --low-mem--trim-adapters --barcode-whitelist --bc-error-thresholds 2 --remove-pcr-duplicates-at-cell-level) (*61*). Only uniquely and properly paired reads with mapping quality greater than 10 were retained using *samtools view* (v1.16.1) with non-default settings (-bhq 10 -f 3) (*65*). Duplicated fragments were removed on a per-barcode basis using *picardtools MarkDuplicates* (v2.27.4) with non-default settings (BARCODE_TAG=CB REMOVE_DUPLICATES=true) (*66*). Alignments were then used to extract single-base resolution Tn5 integration sites by shifting the start and end coordinates by +5 and -4, respectively. Tn5 integration sites were filtered to retain only unique coordinates for each barcode.

#### Variant processing

To partition reads and barcodes by their respective genotypes, we first re-aligned short read whole genome sequencing (WGS) data from all 172 inbred lines (*67*) to the maize B73 v5 reference genome (*60*) using *BWA mem* (v0.7.17) with non-default settings (-M -t 4). Inbred lines with more than one WGS experiment were merged into a single BAM file. WGS alignments for each inbred line were then sorted and deduplicated using *samtools* (v1.16.1) (*65*). Variants were identified using *freebayes* (v1.2.0) with non-default settings (--strict-vcf -n 4 -q 10 --max-coverage 10000) (*68*). To simplify downstream analysis, we only retained bi-allelic SNPs.

Genotype calls with fewer than four reads were set to missing, and variants with more than 25% missing data or a minor allele frequency less than 5% were removed, resulting in a final set of 31,478,930 bi-allelic SNPs for assigning barcode genomes.

#### *In silico* genotype-demultiplexing

To assign barcodes to their genotype(s) of origin, we first removed barcodes with fewer than 100 unique Tn5 integration sites. Genotype and singlet/doublet log likelihoods for individual barcodes were estimated using *souporcell* with non-default values (-k $num_geno --min-alt 5 -- min-ref 5 --skip-remap TRUE --no-umi TRUE) where $num_geno references the number of genotypes multiplexed in the sequencing library (*69*). Barcodes with a greater singlet to doublet log likelihood and the best genotype assignment with 3-fold greater log likelihood than the next best genotype assignment were retained for downstream analysis.

#### Genotyping quality control

To evaluate the reproducibility of chromatin accessibility profiles within a genotype from our approach, we first collected Tn5 insertions sites corresponding to QC-filtered nuclei from a single genotype for each library, independently (representing a single genotype replicate). Next, we counted Tn5 insertion sites intersect with the total set of 108,843 ACRs for each genotype replicate to create a raw counts matrix using *bedtools annotate*. The counts matrix was scaled per million (counts per million) and quantile normalized with the *R* functions, *cpm,* and *normalizeQuantiles* from the *R* packages, *edgeR* and *limma,* respectively. We then estimated Spearman’s correlation among genotype replicates after removing batch effects with the function *removeBatchEffect* from the *R* package, *limma,* with non-default values (batch=library.batch). To determine the correspondence between the variant calls identified by low-coverage WGS and the genotype-aggregates from scATAC-seq, we generate BAM files using QC-filtered nuclei for each genotype. We then identified SNVs using *freebayes* across all 172 BAM files using the same pipeline as described above under the header “Variant processing”. Variant calls from WGS were then merged with variant calls from scATAC-seq using the function *MergeVcfs* from *picardtools* (v2.27.4), removing nonACR-localized SNVs and SNVs on unanchored scaffolds. We then determined the genetic relationship matrix from combined set of variants using the function *snpgdsGRM* from the *R* package, *SNPRelate*, for LD pruned biallelic variants (ld.threshold=0.25 via *snpgdsLDpruning*) with minor allele frequencies greater than 0.05, SNP missingness less than 25%, and method=”Corr”.

#### scATAC-seq quality control

Starting from the unfiltered data set, barcodes with at least 500 unique Tn5 integration sites, standardized (Z-score) fraction of Tn5 integration sites within 2-kb of the TSS greater than the maximum of two standard deviations below the mean and 0.2, and standardized (Z-score) fraction of Tn5 integration sites within ACRs greater than the maximum of two standard deviation below the mean and 0.2, were identified as putative nuclei. To further remove barcodes representing broken nuclei or ambient background, we first used the unfiltered set of barcodes to create a binary sparse matrix of barcodes x 500-bp bins tiled across the maize genome. To determine if a barcode’s genome-wide chromatin profile was indicative of an intact nucleus or background noise, we estimated Spearman’s correlation coefficients for each barcode with the following aggregates normalized by term frequency inverse document frequency (TF-IDF): (*i*) top 1,000 barcodes ranked by Tn5 insertion counts (non-background) and (*ii*) barcodes with less than 100 Tn5 insertions (background). Barcodes with greater correlations to the non-background aggregate relative to the background aggregate were retained.

#### scATAC-seq clustering using reference nuclei

To reduce computational costs and improve cell state identification, we first isolated a reference set of barcodes by removing nuclei from the filtered barcode x 500-bp bin sparse binary matrix with less than 1,000 accessible features using the *cleanData* function of *Socrates* (*56*) with non-default parameters (min.c=1000, min.t=0.0025, max.t=0). As a result, bins with chromatin accessibility frequency of 0.25% or greater across barcodes were retained. The binarized sparse matrix was then transformed using term frequency inverse document frequency (TFIDF) and L2 normalization in *Socrates* using the function *tfidf* with non-default parameters (doL2=T). The dimensionality of TFIDF normalized matrix was reduced to 21 scaled Principal Components (PCs) using Singular Value Decomposition (SVD) using the top 25% of variable features, with the first PC removed due to high correlation (PCC > 0.7) with nuclear read depth via the function *reduceDims* with non-default parameters (method=”SVD”, n.pcs=21, cor.max=0.7, num.var=ceiling(nrow(obj$counts)*0.25), scaleVar=T, doL2=T). Library and genotype effects were removed using *Harmony* (*70*) with non-default parameters (theta=c(2,2), tau=5, sigma=0.1, lambda=c(0.1, 0.1), nclust=50, max.iter.cluster=100, max.iter.harmony=30), return_object=T). The integrated object was used to create a reference embedding using the R package, *symphony* (*71*), via the *buildReferenceFromHarmonyObj* function with non-default parameters (do_umap=F, umap_min_dist=0.01). To visualize chromatin accessibility similarity among reference nuclei, we further reduced the dimensionality of the PC matrix using *umap* with non-default parameters (metric=”cosine”, n_neighbors=30, a=2, b=0.75, ret_model=T). The resulting model was added to the symphony reference object while the UMAP embedding was added to the UMAP slot of the *Socrates* object. Clusters of nuclei with similar chromatin accessibility patterns were identified using the *callClusters* function of *Socrates* with non-default parameters (res=$optres, k.near=30, cleanCluster=F, cl.method=4, e.thresh=3, threshold=3, m.clst=50) using the Leiden graph-based clustering algorithm. The optimum resolution for Leiden clustering was determined by iterating over a range of resolution parameters (0 – 5e^-3^) and estimating clustering stability scores at each resolution. Cluster stability was estimated perturbing 2% of graph edges and taking the clustering consensus values across 5 permutations, as previously described (*72*). The resolution parameter with the greatest cluster stability score was selected for the final clustering call.

To project the remaining cells on to the reference embedding, we first filtered nuclei with fewer than 100 accessible features using the function *cleanData* with non-default parameters (min.c=100, min.t=0, max.t=0). Feature IDs from the query nuclei were then aligned with the reference object to enable downstream integration. The remaining query nuclei were projected into the same TFIDF space as the reference nuclei using the function *projectTFIDF* with default parameters. Nuclei were then split into groups of ∼5,000 and iteratively projected onto the reference SVD embedding. Specifically, during each iteration of new nuclei, we ran *mapQuery2*, a modified scATAC-seq variant of the *mapQuery* from *symphony*, using non-default parameters (svd_diag=obj$PCA_model$d[obj$PCA_model$keep_pcs], vars=c(“library”, “genotype”, sigma=0.01, do_umap=T, scaleVar=F, doL2=T).

#### scATAC-seq cell-type annotation

To annotate cell types, we first estimate gene activity scores by counting unique Tn5 integration sites overlapping gene bodies and in proximal regions 500-bp upstream of TSSs using the *Socrates* function, *estGeneActivity*. Gene activity scores were scaled to sum to 10,000 per nucleus. Cell type annotation was then performed using *a priori* marker gene activity UMAP profiles, cell type marker enrichment, differential gene activity, and multinomial logistic classification as previously described (*56*). Additionally, due to the effects of cell cycle stage towards clustering and interpreting cell type annotations, we first collected a list of known cell cycle stage-specific genes (*73*). We then estimated the enrichment of genes specific to each stage (average normalized gene activity) versus 1,000 permutations of the same number of randomly selected cell cycle genes within each nucleus. Final cell cycle stage scores (Z-scores) were derived by standardizing the average normalized gene accessibility of the focal stage by subtracting and dividing the mean and standard deviation of the permuted enrichment scores, respectively. The cell cycle stage with the highest normalized score was used as the predicted cell cycle stage. Final cell state annotations were generated by considering collection of predicted cell type and cycle stage information.

#### ACR identification

A comprehensive set of ACRs were first identified using aggregated Tn5 insertion counts for each genotype x cell type combination using MACS2 (v2.2.7.1) (*74*) with non-default parameters (-g 1.6e9 --nomodel --keep-dup all --extsize 150 --shift -75 --qvalue 0.05). ACRs from all genotype by cell-type combinations were then concatenated into a single master list (n=65,450,651). To address the presence of overlapping intervals without merging, we recursively selected the most significant ACR (based on -log_10_ q-value) and removed all ACRs with lesser -log_10_ q-values and direct overlap with the focal ACR, repeating the process iteratively to produce a final set of non-overlapping ACRs (n=4,323,286). For each cell type, we retained ACRs from the non-overlapping set that were supported by ACR calls (intersection >= 1bp) from at least two distinct genotypes, resulting in 631,846 ACRs. To remove ACRs with chromatin accessibility enrichment similar to the background Tn5 insertion rate, we collected the same number of random genomic regions (n=631,846) matched by interval size specifically from mappable regions. Mappable regions were defined as genomic regions covered by at least one read from a simulated data set generated by *wgsim* (*65*) with non-default parameters (-N 5e8 -1 100 -2 100 -d 300). We then constructed binary sparse matrices for (1) control regions x cell and (2) ACRs x cell and estimated the average Tn5 insertion frequency for each feature across all cells in each matrix. The average Tn5 insertion frequency across cells for each control region was then used to fit a beta distribution with *fitdist* from the *fitdistrplus* R package. We obtained *P*-values for each ACR by comparing the observed Tn5 insertion frequency to the null model parameters derived from control regions using the base R function *pbeta*. *P*-values were corrected for multiple testing using *p.adjust* with non-default parameters (method=”fdr”).

ACRs with an FDR less than 0.01 were retained, resulting in a total of 108,843 ACRs. For analyses specific to *cis*-regulatory function, we estimated the proportion of each ACR corresponding to exonic, intronic, intergenic (>2-kb upstream of the TSS), 50-bp surrounding the TSS, and proximal (<2-kb upstream of the TSS) sequences. We then removed all ACRs where the majority of the ACR interval overlapped an exon, except in cases where an ACR also overlapped with a TSS. Altogether, we identified a total of 82,098 ACRs with putative *cis*- regulatory function for downstream analysis. ACRs were defined as conserved if at least 75% of genotypes exhibited at least 90% shared sequence alignment with B73. Cell-state specificity was estimated via the Tau metric (*75*) and ACRs with Tau greater than 0.7 were considered as cell-state-specific.

#### snRNA-seq raw data processing

We re-analyzed one previously generated snRNA-seq dataset (NCBI GEO accession number: GSM5344025) from the same tissue type in conjunction with the three newly generated snRNA-seq data (*56*). For the new data sets, binary base call sequences files (BCL) output from the Illumina S4 NovaSeq 6000 were demultiplexed with *cellranger mkfastq* (v6.1.1). Now using all new and previously generated data sets, raw reads were mapped with *cellranger count* resulting in BAM files containing all reads (both mapped and unmapped) with whitelist-corrected barcodes in the CB tag and UMI sequences in the UM tag. Alignment files were then filtered to remove low mapping quality alignments with *samtools view* (v1.16.1) (*-q 30*) and to remove duplicate reads using the *picardtools* (v2.27.4) function, *MarkDuplicates* by setting the BARCODE_TAG parameter equal to the concatenation of the CB and UMI sequences residing in a custom tag labeled UM. We then converted the filtered BAM alignments into BED-formatted files by moving the cell index and UMI sequence to the read name and piping the output to *bedtools bamtobed* (v2.27.1). Read counts per gene per cell were estimated by counting the overlap of alignments with gene bodies (including introns) for each gene via a custom perl script. Barcodes with fewer than 750 UMIs and 250 transcribed genes, and greater than 10,000 transcribed genes were removed. Barcodes with more than 5% of UMIs mapping to chloroplast or mitochondrial genes were also discarded, retaining barcodes where 95% of UMI were derived from the nuclear genome. We also filtered cell barcodes with gene expression patterns more correlated to background transcript profiles, similar to scATAC-seq data processing (see above). Cell genotypes were identified using *souporcell* (*76*) with non-default parameters (-- threads 30 -k 2 --known_genotypes $vcf --known_genotypes_sample_names Mo17 B73), with barcodes identified as doublets removed from the analysis.

#### Multimodal data integration and cell-type annotation

Gene chromatin accessibility and transcript abundance matrices were first filtered to retain genes accessible/transcribed in at least 0.1% of nuclei (and nuclei with at least 100 accessible/transcribed genes) and matching between modalities. The ATAC and RNA matrices were loaded into the R package, *rliger* (*77*), and normalized with the function *normalize* with default parameters. The top most variable genes were selected using the function *selectGenes* with non-default parameters (datasets.use=”RNA”, var.thresh=0.5) and the matrices from both modalities were scaled without centering using the function *scaleNotCenter* to preserve the non-negative distributions expected by non-negative matrix factorization. Joint non-negative matrix factorization was performed with the function *optimizeALS* with non-default parameters (k=20, lambda=5). Finally, we performed quantile normalization using *quantile_norm* with default parameters to fully integrate the cell factor loadings from the two modalities. Nuclei clustering was performed similarly to the scATAC-seq data with the exception of setting k=15 and using 20 cell factors loadings returned by quantile normalization when calling clusters with the consensus-based approach. Cell type annotation was performed identically as described for the scATAC-seq only data set, including visualization of marker genes via UMAP and cluster-aggregated heatmaps, estimation of differentially accessible and transcribed genes, and automated per-cell annotations via multinomial regression.

#### phyloP score generation

Generation of phyloP scores was done in a multi-step process. Genome alignments for 12 monocot genomes, including *Zea mays, Sorghum bicolor, Oryza sativa, Brachypodium distachyon, Urochloa fusca, Panicum halli, Hordeum vulgare, Panicum miliaceum, Setaria itatlica, Eleusine coracana, Saccharum spontaneum, Miscanthus sinensis* was done using progressive *cactus* (version 2.6.7) (*78*) on softmasked genomes via RepeatMasker. After whole genome alignment, neutral sites were identified in the *Zea mays* genome to act as a null model. To do so, functional genomic data in the form of ChIP-seq peaks associated with functional markers correlated with transcription (H3K36me3, H3K4me1, H3K4me3, and H3K56ac from (*19*)) were removed. Additionally, regions of the genome with evidence of transcription (greater than 5 RNA-seq reads from reference (*79*) aligning to any given locus), as well as regions overlapping exons and introns were removed. All neutral sites were required to additionally be found in at least 9 of the provided genomes.

The isolated neutral sites were then aggregated together and run through *phyloFit* (version 1.5) (*80*) with the default options including ‘--subst-mod REV --msa-format MAF’. The resulting model “mod” file as well as the location of the neutral sites were then used as input for the hal implementation of phylop, halTreePhyloP.py version 2.0 with defaults (*81*). The resulting bedgraph files were used for downstream analysis.

#### ACR conservation analysis

To determine the extent of sequence conservation in ACRs across unique inbred maize lines and the wild progenitor, teosinte, we estimated the fraction of each ACR covered by at least one read after merging available whole genome sequencing data (Goodman-Buckler diversity panel (*67*) and twenty teosinte (*82*) genotypes), 150bp simulated reads derived from available reference assemblies (NAM founders (*60*) and TIL11 (*83*)), and the present scATAC-seq data. Simulated paired-end reads were constructed using the default parameters of *wgsim* from the *samtools* suite, setting read lengths and total paired read counts to 150bp and 500,000,000, respectively. Domesticated maize-specific ACRs were identified as those lacking alignments (< 10% ACR coverage) across all 21 teosinte genotypes. For the Monte Carlo permutation ridge regression test, we scored the presence/absence of a given TFBS within an ACR for each genotype based on complete overlap at least one sequencing read from the merged alignments. Ridge regression models were fit (ACR conservation score ∼ proportion of genotypes with TFBS present) with *cv.glmnet* from the *R* package, *glmnet* with non-default parameters (alpha=0). The observed beta values were compared to null beta distributions from 1,000 permutations of randomizing the row and column IDs. Z-scores were determined as the difference between the observed beta and mean null distribution, divided by the standard deviation of the null beta distribution. *P*-values were computed by comparing the observed beta with the null distributions. FDR was estimated using the *R* function, *p.adjust* with non-default parameters (method=”fdr”).

#### Preprocessing for caQTL mapping

Prior to running caQTL analysis, we first restricted the ACR set to regions where genotype and/or cell type was significant (FDR < 0.05) in a generalized logistic regression model that included (a) log_10_-transformed number of accessible sites, (b) sequencing library ID, (c) genotype, and (d) cell cycle stage. For identification of cell-type-associated ACRs, we also included the first five principal components from PCA of 31M bi-allelic SNPs. For identification of genotype-associated ACRs, we included the first two components from the symphony embedding. A total of 57,984 non-exonic ACRs varied significantly among genotypes, cell types, or both were identified as candidates for caQTL mapping. To estimate the relative variance explained by cell type and genotype, we ran *fitExtractVarPartModel* from the R package, *variancePartition* using cell type and genotype as random effects and all covariates as fixed effects.

To enable efficient caQTL mapping, we adopted a pseudobulking approach by aggregating Tn5 insertions of cells corresponding to the same cell type and genotype across all ACRs. The raw counts matrix was scaled by counts per million (CPM), quantile normalized, and subjected to inverse normalization per ACR to mitigate the effects of outliers. After partitioning the *genotype-cell type* (columns) x *ACR* (rows) matrix by cell type, we ran PCA for each distinct cell type matrix composed of *genotype* x *ACR* with CPM-scaled and ACR-standardized counts.

Simultaneously, we estimated the top 10 PCs from the 31,478,930 bi-allelic SNVs used to genotype individual cells using the *R* package, *SNPRelate* (*84*). For each *cell type* x *genotype* combination, we also collected information on the total number of Tn5 insertions within ACRs, the total number of cells, the fraction of cells derived from all sequencing libraries, and the fraction of cells corresponding to G1, G1/S, S, G2, or M phases of the cell cycle. Technical covariates were subjected to PCA to identify orthogonal sources of variation, with the top X PCs explaining 95% of the variation (depending on cell type) merged with the first 20 cell-type pseudobulk chromatin accessibility PCs and first five genetic PCs.

#### caQTL mapping

For identification of cis*-*associated genetic variants in bulk tissue and each individual cell type, we used *tensorQTL* (*62*) with a max cis distance of 1 Mb. To maintain consistency with prior GWAS and eQTL studies in maize, bi-allelic SNV calls were collected from the maize HapMap3 and uplifted to AGPv5 (*67*). After removing inbreds lacking scATAC-seq data (*n* retained = 172), we filtered the bi-allelic SNVs to remove sites with MAF less than 0.01 and missingness greater than 90%. Normalized phred genotype likelihoods were used to determine allele dosages. All missing values were then imputed using the nearest neighbor via the *R* package, *impute* (*85*), via the function *impute.knn* with non-default parameters (k=10) for overlapping windows of 10,000 variants (step size = 5000) and each chromosome independently. We then removed variants with less than 8 alternate alleles across the 172 individuals (rowSums(x) > 8). Finally, the imputed SNV set was converted to *plink2* (*86*) format with non-default parameters (--make-pgen --output-chr chrM --vcf $VCF dosage=DS). To identify caQTL, we ran *tensorQTL* with non-default parameters (--mode cis --covariates $cov --qvalue_lamba 0.85 --dosage). To retrieve all nominal *P*-values for downstream analysis, we ran *tensorQTL* with non-default parameters (-- covariates $cov --dosages --mode cis_independent). We set a dynamic FDR (between 0.01 and 0.1) across cell types based on a linear model fit to the average Tn5 integration depth per genotype, where more stringent FDR thresholds were set for cell types with more sequencing information.

Identification of eQTL was performed similarly as caQTL with the exception of independent construction of covariates, which was conducted identically as the scATAC-seq data. Briefly, we aggregated the top 20 PCs from the standardized genotype x gene matrix with the top five genetic PCs based on the SNV genotypes from *SNPRelate*. The log_2_ CPM normalized genotype x gene transcript abundance matrix was obtained from a prior study (*31*). Gene IDs were converted to v5 names using the liftover function of *CrossMap*. Gene expression was inverse quantile normalized per gene to mitigate the effects of outliers prior to eQTL mapping. We used identical functions and parameters of *tensorQTL* to identify eQTL as described for caQTL above.

#### Identification of cell-state-specific effects

Variant-level effect size and standard error summary statistics from *tensorQTL* for all ACR by SNV tests in each cell state were extracted and merged into two independent matrices. These matrices were then loaded into the *R* package, *mashr* (*34*), with the function *mash_set_data.* We estimated the data-driven covariance matrices using the top five PCs using the most significant (lowest *p-*value) caQTL for each caQTL-ACR with the function *cov_pca* with non- default settings (npcs=5, subset=topQTL), followed by extreme deconvolution with the function *cov_ed* with non-default parameters (subset=topQTL). We also determined the canonical covariance matrices using the function, *cov_canonical,* with default settings. The joint data-driven and canonical covariance matrices were used to fit the multivariate adaptive shrinkage model with the function, *mash*. Pairwise sharing of the caQTL among cell states was determined by the function *get_pairwise_sharing* with default settings using only the top caQTL per ACR as input. Individual caQTL for each cell state was considered shared if the absolute posterior effect size was within a factor of two of the absolute maximum effect size across cell states and the same sign as the most significant cell state context.

#### Fine-mapping caQTL

Statistical fine-mapping of caQTL was performed using *susieR* (*87*). Briefly, we first identified all SNVs within 100bp of caQTL-ACR coordinates and extracted the caQTL summary statistics output from *tensorQTL*. We then ran *susie_rss* with non-default parameters including the SNV correlation matrix adjusted by the same covariates given to *tensorQTL,* the beta hat and beta hat standard error from *tensorQTL* summary statistics, and set *L*=10 and *estimate_residual_variance*=T. Finally, we kept all SNVs within the 95% credible set list (the list of variants with 95% probability that the causal variant is included) with posterior inclusion probabilities greater than zero and FDR values less than the dynamic threshold described above.

#### Differential allelic TF footprinting

To identify TF footprints differing between alleles, we identified all SNVs within and flanking ACRs up to 1-kb away. We then identified the top two most frequent haplotypes of each ACR by clustering genotypes by their SNV dosage profiles using the R package, *mclust* with non-default parameters (G=1:10). Genotype membership for each haplotype was recorded across all ACRs, retaining ACRs with at least 5 individuals represented in the top two most frequent haplotype groups. Then, we aggregated scATAC-seq alignments for each ACR and each cell type to create pseudobulk aggregates reflecting the top two distinct haplotypes at cellular resolution.

These alignments were then used as input for *TOBIAS* (*88*) along with custom reference FASTAs generated by *FastaAlternateReferenceMaker* from the *GATK* suite of tools for each haplotype. *TOBIAS* was run with default parameters for the subfunctions, *ATACorrect* and *ScoreBigwig. TOBIAS BINDetect* was run with non-default parameters to increase the stringency in motif identification and footprint calling (--motif-pvalue 1e-5 --bound-pvalue 1e-4).

We set an additional filter where differential TF footprints were then identified as regions with footprint scores greater than 5 in at least one haplotype and “unbound” based on *TOBIAS* thresholds in the other haplotype (footprint score bound/unbound threshold range across cell types = 0.95 - 1.58).

#### TF motif variant effects

Comparison of alleles overlapping predicted TFBSs while considering TF motif position weight matrices has been previously used to predict allelic effects on TF binding affinity (*89*). To disentangle the potential mechanisms of chromatin accessibility variation, we ran *motifbreakR,* an R package designed to estimate allelic effects on TF binding affinity (*89*). Specifically, we loaded the same variants used for caQTL mapping reformatted in BED as required by *motifbreakR,* as well as the coordinates of all caQTL-ACRs from cell state-resolved caQTL mapping. The SNV list was filtered to retain records embedded within the test set of ACR coordinates. We then ran *motifbreakR* using all plant-derived JASPAR 2022 motifs from the *MotifDb* R package with non-default parameters (filter=T, threshold=1e-5, method=”ic”, show.neutral=T).

#### Fine-mapping and TF footprint STARR-seq validation

To estimate enhancer activity of each allele for TF footprints containing fine-mapped variants, we first collected the reference and alternate sequences of all TF footprints containing fine-mapped variants. We then identified all reference and alternate allele TF footprint sequence occurrences in the genome for regions containing at least one mapped DNA or RNA STARR-seq read using *seqkit locate* (v2.5.1). Enhancer activity was estimated as previous described (*90*), which was used to map (via *bedtools map*) the maximum enhancer activity onto all reference and alternate TF footprint sequence occurrences. The enhancer activity of a given fine-mapped TF footprint allele was taken as the average enhancer activity across all occurrences genome wide, requiring at least one occurrence for both reference and alternate alleles for a given fine-mapped TF footprint. We then estimated the allelic effect as the difference in averaged enhancer activity between alleles. For fine-mapped TF footprints with multiple genome wide occurrences for each allele (16,646 fine-mapped TF footprints), we further determined if alleles exhibited significant differences in enhancer activity using the Wilcoxon Rank Sum Test and correcting the nominal *P*-value using the Benjamini-Hochberg correction and kept records with FDR < 0.05.

#### Construction of cell-state-specific gene regulatory networks

Since GRN inference benefits from non-sparse data, we first constructed metacells using the snRNA-seq and scATAC-seq integrated embedding. Specifically, we identified the reciprocal 10 nearest neighbors between snRNA-seq and scATAC-seq nuclei using the cell factor loadings from iNMF via the function *get.knnx* from the *R* package, *FNN.* The imputed transcriptome and chromatin accessibility profiles for scATAC-seq and snRNA-seq nuclei, respectively, was then calculated by taking the mean across the 10 nearest neighbors from the other modality. To create metacells, we first identified metacell seeds by randomly sampling 10% of nuclei from each cell state. The 50 nearest neighbors from the integrated embedding to metacell seeds were determined using *get.knn,* with nuclei assigned to multiple metacell seeds randomly removed such that a single nucleus was assigned to only a single metacell seed. We then determined the chromatin accessibility and transcriptome profiles of each metacell as the average of all nuclei assigned to the seed. ACR chromatin accessibility scores for each metacell reflected the fraction of assigned nuclei with at least one Tn5 insertion site, whereas metacell gene transcript abundance represented the average log-normalized read counts. Finally, we scaled gene transcript abundances such that the total normalized counts across all genes of a metacell summed to 100,000.

To identify GRNs for each cell state, we first loaded the normalized paired transcriptome and chromatin accessibility counts for each cell state into Seurat (v5) using *CreateSeuratObject* and *CreateChromatinAssay* functions. Maize TF motifs were collected from the JASPAR2022 database, and maize-*Arabidopsis* TF orthologs were collected from a previous study to infer DNA binding preferences of maize TFs lacking experimental data (*56*). All downstream GRN construction was conducted using the *R* package, *Pando* (*91*). The GRN was then initiated by the function *initiate_grn* with default parameters. TF motifs within ACRs were identified using the function *find_motifs* using the aggregated maize JASPAR2022 motifs from above as input. TF effects on gene expression of other TFs was then determined using the function *infer_grn* with non-default parameters (genes=TF.ids). TF modules were identified using the function *find_modules* with more stringent parameters (p_thresh=0.05, rsq_thresh=0.1). The normalized TF effect score was determined as previously described (*91*). Associations with FDR < 0.05 were retained in the final data set.

#### Co-accessibility QTL mapping

To identify genetic variants putatively affecting higher-order chromatin interactions, we first estimated co-accessibility on a per-genotype basis. Briefly, we collected all cells (*n*) from each genotype in isolation, constructed pseudocells with the k-nearest neighbors (where *k* is the square root of *n*) and scaled Tn5 integration counts per ACR by counts per million. Raw co-accessibility scores (*A*) between all ACRs < 500-kb and > than 2-kb apart were then estimated using Spearman’s correlation coefficient. We then penalized raw co-accessibility scores by the distance between ACRs to reflect the exponential decay observed empirically between pairs of interacting genomic regions using equation (Eq.) 1:

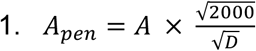

Where *D* is the distance between the pair of ACRs. Finally, we estimated residuals between the penalized co-accessibility scores and a null set of permuted penalized co-accessibility scores derived from randomizing the cell x ACR matrix that maintained the same number of accessible cells per ACR. The set of co-accessible ACRs found across all genotypes was then filtered to retain links supported by previously published maize HiC and HiChIP chromatin loops (*19*).

Genetic variants greater than 5Mb away from the focal co-accessible ACRs were then identified using *tensorQTL* with non-default parameters (--mode trans --covariates $cov --qvalue_lamba 0.85 --dosages --pval_threshold 0.00001 --batch_size 1000). Because transQTL mapping results in a substantially larger test space, we opted to focus on the number of traits associated with a single variant at *P<*1e^-5^ rather than using multiple test correction. Candidate genes were identified by first selecting all *trans*-QTL variants associated with more than 30,000 co-accessible ACRs and collecting gene IDs within 50-kb of the top variants.

#### Long-range caQTL analysis

To identify caQTL that potentially influence accessibility at other loci, we first identified caQTL- ACRs containing local caQTL (caQTL within the ACR coordinates). We then kept all caQTL records that were associated with two or more ACRs. The set of randomly permuted ACR-ACR interaction was derived by randomly linking ACRs from the long-range caQTL-ACR data set (i.e. scrambling the association between two ACRs using the same set of observed ACRs).

Predicted interactions were determined by entering the long-range caQTL-ACR coordinates in BEDPE into the GenomicLinks web browser (*41*).

To assay the distribution of TF motifs among co-accessible ACRs with the support of caQTL evidence, we organize co-accessible ACRs into two main groups for analysis. The first group, our test set, includes co-accessible ACRs that have at least one caQTL and evidence from HiC/HiChIP experiments (*19*) showing they physically interact. The second group, our control set, consists of co-accessible ACRs showing physical interactions in HiC/HiChIP data but lacking caQTL support. To evaluate the enrichment of specific TF motifs within these groups, we employed the Fisher’s Exact Test for Count Data in R (function: fisher.test, alternative=“greater”, R version 4.3.3). The contingency table was defined as follows: *matrix(c(n_i_, d_i_, c_i_, t_i_), nrow = 2)*. Here, ’*n_i_*’ represents the total number of co-accessible ACRs with a TF motif overlap, excluding those supported by both caQTL and HiC/HiChIP data; ’*d_i_’* denotes co-accessible ACRs with HiC/HiChIP evidence but lacking caQTLs where the TF motif *i* is present; *’c_i_’* accounts for co-accessible ACRs supported by caQTL but not HiC/HiChIP; and ’*t_i_*’ capture co-accessible ACRs supported by both caQTL and HiC/HiChIP data where the TF motif *i* is found. The detection of TF motifs in co-accessible ACRs was performed by scanning the corresponding PWM across all co-accessible ACRs. We only counted hits where the PWM was found in both anchors of the co-accessible ACR. The scanning of TF motifs was carried out with the *Motifmatchr* package in R (v1.24.0) with default settings. To minimize the redundancy of TF motifs arising from different experiments of the same TF or from TFs within the same family with similar TF motifs, we aligned all PWMs and extracted their PWM signatures using the *clusterMotifs* and *motifSignature* functions (with parameters: cutoffPval < 0.0001 and min.freq = 1) from the *R* package *motifStack* (v1.46.0) (*92*). Finally, all PWM were collected from JASPAR 2024 keeping only PWM reported for *Arabidopsis thaliana* (*93*).

#### Complex trait association analyses

Forty two agronomic traits of maize were downloaded from https://www.panzea.org/phenotypes (full 277 genotypes) and SAM morphology traits (172 genotypes matched to scATAC-seq data) were acquired from (*94*). We calculated best linear unbiased estimates (BLUEs) from phenotypes collected in multiple years or environments (genotype treated as fixed effects and environment/location treated as a random effects) using the *R* package, *lme4*. Direct trait estimates were used in the absence of environment and/or year replication. The genome-wide association analysis (GWAS) was conducted using the entire Goodman-Buckler diversity panel (*n*=277) for a subset of SNVs within 500-bp of any ACR (*n*= 2,574,564) starting from the same SNV set as for caQTL mapping. For the agronomic traits (where *n* genotypes = 277), we used a fixed effects generalized linear model approach, implemented in the *R*-based software, *GAPIT v3* (*95*). The first five genetic PCs were included in the model to correct the hidden population structure. The genome-wide significance threshold was determined by pruning the 2.5M SNVs by LD with plink2. Specifically, we removed variants with the --indep-pairwise function with a 5- kb window size and r^2^ threshold of 0.2. These parameters produced a filtered set of 493,100 independent variants, resulting in a Bonferroni threshold of -log_10_(*P*-value) of 5.69 [- log_10_(1/493100)]. All GWAS records with *P-*values below the Bonferroni threshold or Benjamini-Hochberg FDR < 0.05 were retained. Finally, we retained significant GWAS hits either identical or in LD (r^2^ > 0.5) with previously identified quantitative trait nucleotides (QTNs) for the same traits (*2*).

To estimate the phenotypic variance (for SAM morphology and agronomic traits) explained by each SNV, ANOVA was used with contrasting linear models Eq. 2 and Eq. 3:

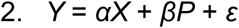

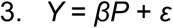

Where *Y* is the phenotype, *X* is the SNV genotype, *P* is the PCs, *α* is the SNV effect, *β* is the PC effects, and *ɛ* is random error, which follows a N(0, σ^2^_e_) distribution. Thus, the variance due to each SNV was reported after adjusting for the population structure effects, as previously described (*96*). We estimated the significance of the proportion of variance explained (PVE) of caQTL versus non-caQTL by first partitioning caQTL by cell state and randomly selecting the same number of non-caQTL SNVs. We compared the mean SNV PVE for caQTL and non-caQTL SNVs across 10,000 permutations.

For the transcriptome-wide association study (TWAS) and chromatin accessibility-wide association study (CAWAS) across all measured traits in the 172 genotypes, we re-distributed the normalized bulk CPM value for each gene and each ACR into a consecutively numeric range from 0 to 2, with the implementation of mixed linear model (MLM) in the *R* package, *rMVP* (*97*), using a relatedness matrix based on gene expression and ACR coverage intensity, respectively, to correct for population structure. Multiple testing corrections were performed with a false discovery rate (FDR) of 0.05 using the R function *p.adjust* (method = “fdr”). The relative contribution of individual TF motifs towards phenotypes was determined by matrix multiplication of the CAWAS results with a binarized motif x ACR matrix, scaling the subsequent motif scores from 0 to 1 across all motifs for each trait.

#### Colocalization analysis

Colocalization analysis for GWAS and caQTL was performed using the SuSiE workflow of *R* package, *coloc* (v5.2.3) (*98*). Because colocalization requires dense summary statistics, we reran GWA for all complex traits using the full set of SNVs across the 277 inbred individuals. Z-scores derived from GWAS and caQTL in a 1Mb window surrounding each caQTL-ACR, along with LD matrices, were used as inputs. For each cell state and GWAS combination, the function *runsusie* was run with non-default parameters (L=10, coverage=0.9). In a handful of cases where *runsusie* failed to recover fine-mapped variants, we used the default Approximate Bayes Factors approach of *coloc*. Variants with posterior probability of colocalization (PP.H4) ≥ 0.8 and previously identified as caQTL (see section “caQTL mapping” above) were considered as colocalized variants.

#### Linkage disequilibrium analysis

To identify a confident set of molecular QTL associated with trait variation, we first selected caQTL variants where the same SNP was also an eQTL. We then estimated the pairwise linkage disequilibrium (LD) between shared molecular QTL (caQTL + eQTL) and all GWAS hits within 10-kb using the following formula in Eq. 4

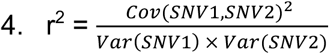

Here, we considered haplotype signals between the molecular and complex traits to be shared at linkage disequilibrium r^2^ > 0.8 (i.e. LD_caQTL-eQTL_ = 1 and LD_caQTL-GWAS_ > 0.8) simultaneously.

#### Analysis of population differentiation and adaptation

We estimate population differentiation at single-nucleotide resolution as Weir and Cockerham’s *F*_ST_ using *vcftools* (v0.1.16) with non-default settings (--weir-fst-pop temperate_genotypes.txt -- weir-fst-pop tropical_genotypes.txt) where we randomly sampled temperate lines to have the same number of individuals in each population (*99*). Chromatin accessibility QTL SNVs were split into low and high *F*_ST_ groups using the R function*, ntile*, from the package *dplyr* using non-default settings (caQTL$fst, n=2). Subpopulation-specific ACRs were identified as ACRs with FDR<0.1 from multiple-tested corrected *P-*values using *p.adjust* (method=”fdr”) output from a Wilcoxon rank sum test using the inverse-normalized pseudobulk chromatin accessibility counts (identical to the input used for caQTL mapping). The FDR<0.1 threshold was biologically motivated by the locus, *Vgt1* (FDR=0.057), that is known to regulated differentially between temperate and tropical inbreds. ACRs under selection were identified by intersection (bedtools intersect) with a previously generated list of genomic regions under selection (*47*).

## Supplementary Text

### Justification of ACR prefiltering

As stated under the header “Preprocessing for caQTL mapping” in the Materials and Methods, we selected ACRs with significant genotype and/or cell type effects using a generalized logistic regression model. Under typical conditions, pre-filtering for features correlated with the probability of finding a significant effect (termed dependent filtering) has been shown to increase FDR (*100*). However, since our caQTL calling pipeline included consideration of variants across the genome (unrestricted by co-localization with ACRs; ∼31M SNVs), we reasoned that the increased multiple testing burden would offset FDR inflation due to dependent filtering. To test this, we performed caQTL mapping on the bulk tissue data with the full set of 82,098 ACRs and a reduced set of 481,947 ACR-localized variants (independent filtering), revealing a total of 37,043 caQTL-ACRs (compared to our reported value of 23,959). The independent filtering approach recovered 97.3% of caQTL-ACRs (23,315/23,959) defined by our approach outlined in the Materials and Methods. Therefore, we conclude that caQTL workflow outlined in the Materials and Methods is robust to FDR inflation due to dependent pre-filtering.

**Fig. S1:**
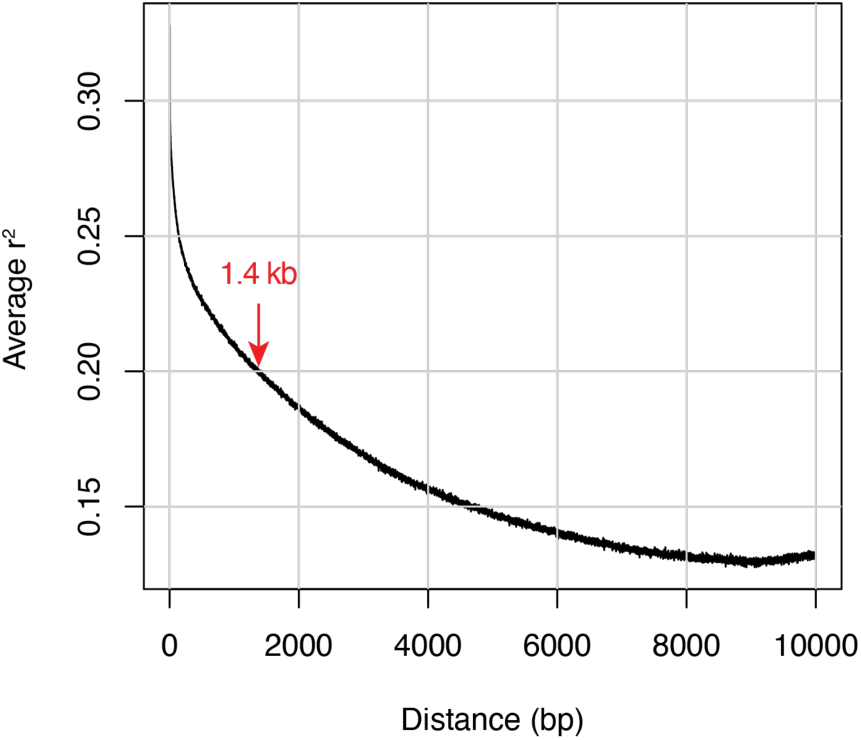
**Genome-wide average linkage disequilibrium decay in the Goodman-Buckler diversity panel in the present study (n=172) as a function of physical distance.**

**Fig. S2:**
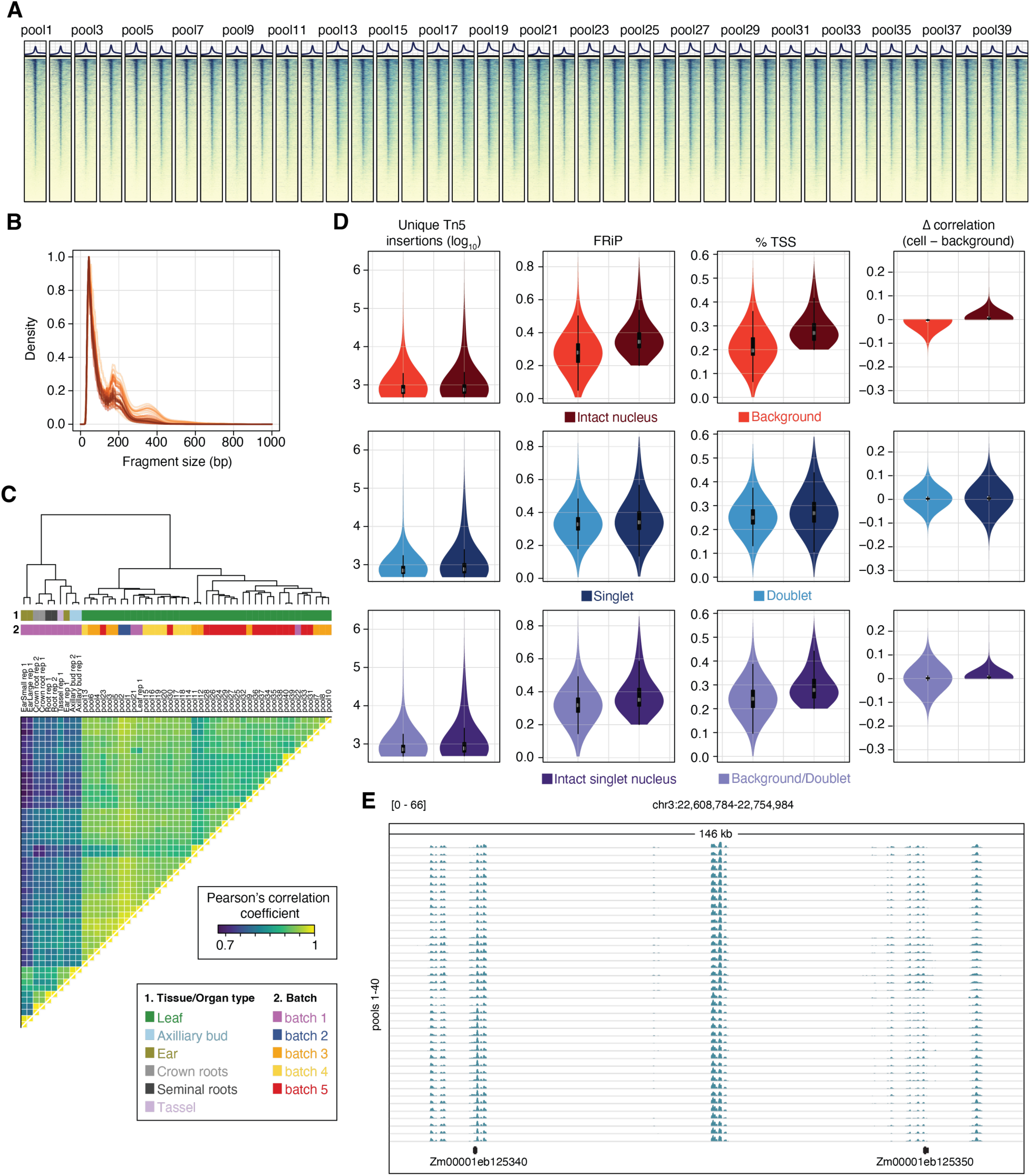
Quality control analysis. (**A**) Meta plot and heatmap of Tn5 insertions sites surrounding all ACRs across 40 genotype-pooled scATAC-seq libraries. (**B**) Fragment size distributions across all 40 genotype-pooled scATAC-seq libraries. (**C**) Correlation analysis from the union of all ACRs among the 40 seedling genotype-pooled libraries and previously published maize scATAC-seq libraries. (**D**) Barcode quality control filtering metrics. (**E**) Representative genome browser view of nuclei-aggregated normalized read depths across all 40 genotype-pooled scATAC-seq libraries. Row tracks are ordered from pool1 to pool 40.

**Fig. S3:**
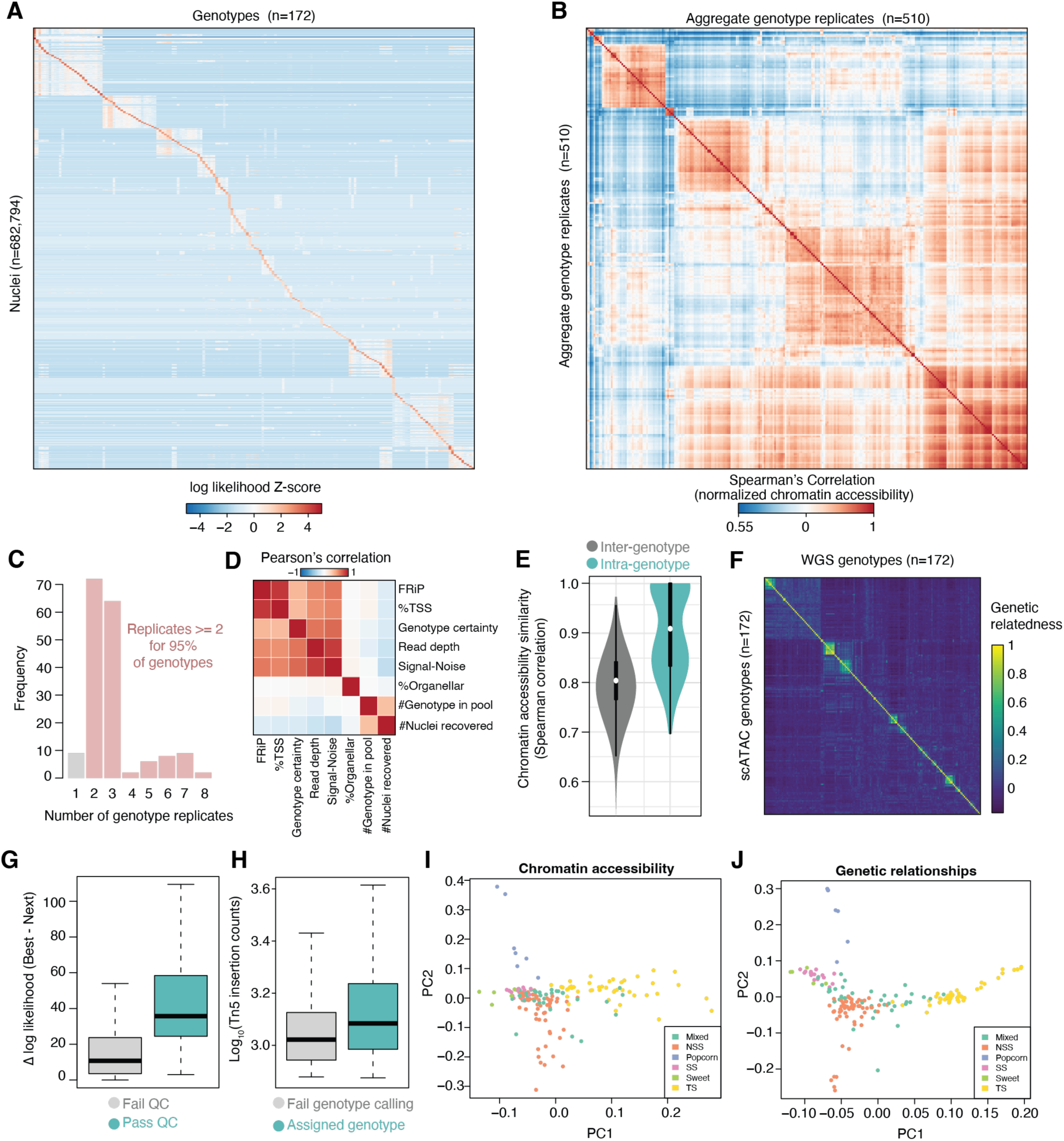
Nuclei genotyping quality metrics. (**A**) Standardized genotype assignment log likelihoods for each barcode (rows) and possible genotype (columns). (**B**) Pairwise correlations among aggregated genotype chromatin accessibility profiles split by replicate. (**C**) Frequency of genotype replication. (**D**) Correlation among various quality metrics and pooling parameters (# of genotypes in a pool and # of nuclei recovered). (**E**) Comparison of chromatin accessibility profiles between genotypes (inter-genotyping) or between genotype replicates (intra-genotype). (**F**) Genetic relationship matrix between scATAC-seq variants and WGS-based variants. Genotype ordering is identical in the rows and columns. (**G**) Difference in log likelihoods between the best and next best genotype assignments split by QC status. (**H**) Distribution of Tn5 insertions counts per barcode for nuclei passing or failing genotype calling. (**I**) Scatter plot of the first two PCs based on standardized chromatin accessibility values of nuclei-aggregates grouped by genotype. Points are colored by subpopulation assignment. (**J**) Scatter plot of the first two PCs based on SNVs identified within scATAC-seq nuclei-aggregates grouped by genotype. Points are colored by subpopulation assignment.

**Fig. S4:**
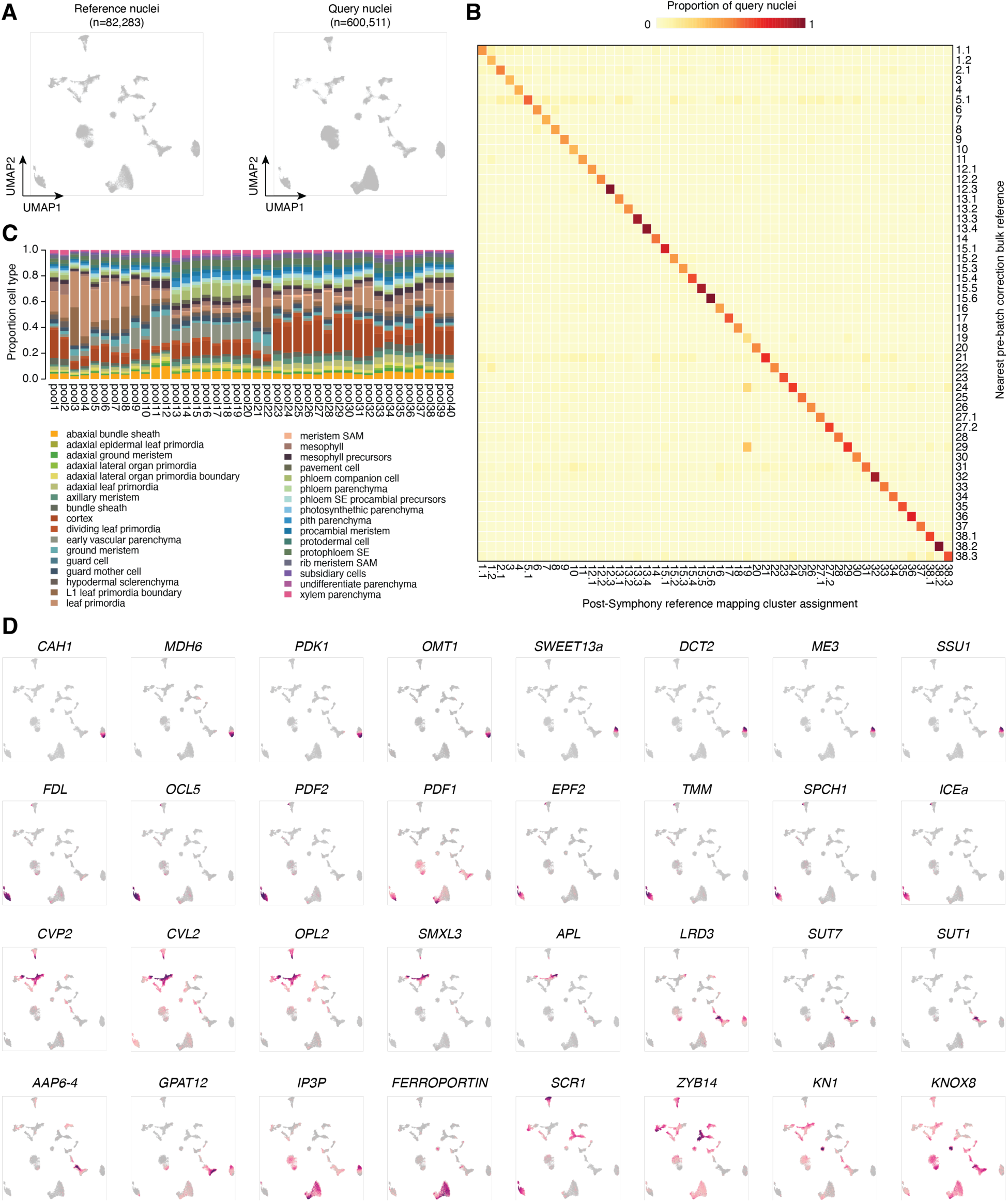
Reference embedding construction. (**A**) Comparison of UMAP embeddings of reference (left) and query (right) nuclei. (**B**) Heatmap of the proportion of query nuclei with the same nearest cluster assignment (based on Spearman’s correlation to the cluster bulk) pre- and post-reference mapping. (**C**) Cell state proportions across each genotype pool. (**D**) UMAP embeddings colored by marker gene chromatin accessibility scores.

**Fig. S5:**
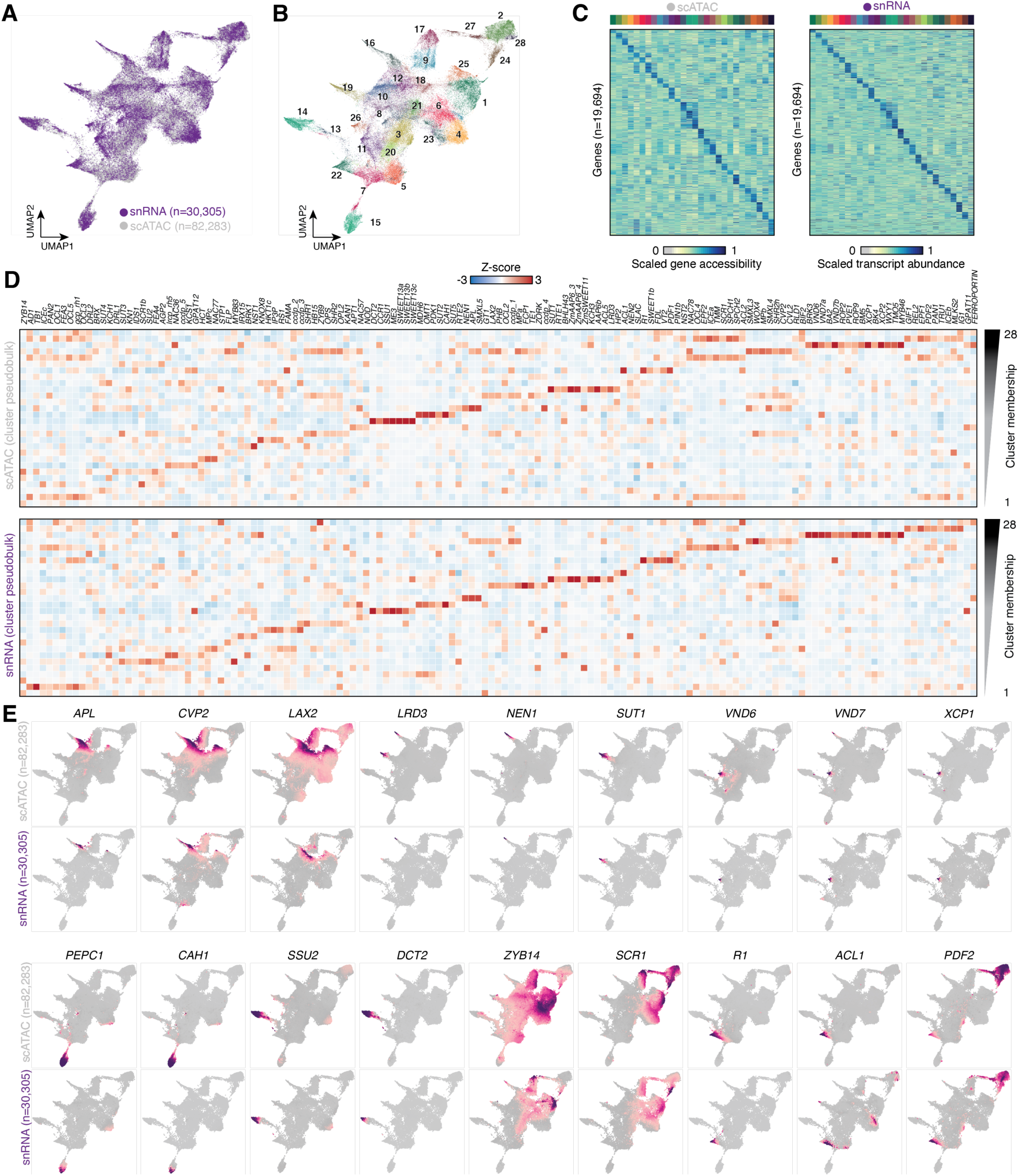
Integration of seedling nuclei scATAC-seq and snRNA-seq. (**A**) UMAP embedding of maize seedling snRNA-seq (purple) and reference scATAC-seq nuclei (grey). (**B**) UMAP embedding of leiden clusters of the integrated embedding. Nuclei are colored by cluster assignments. (**C**) Comparison of scaled gene chromatin accessibility (left) and transcript abundances (right) for cluster aggregates across 19,694 expressed genes. (**D**) Marker gene chromatin accessibility (top) and transcript abundances across clusters identified in the integrated embedding. (**E**) UMAP embedding colored by marker gene chromatin accessibility and transcript abundances for scATAC-seq and snRNA-seq nuclei, respectively.

**Fig. S6:**
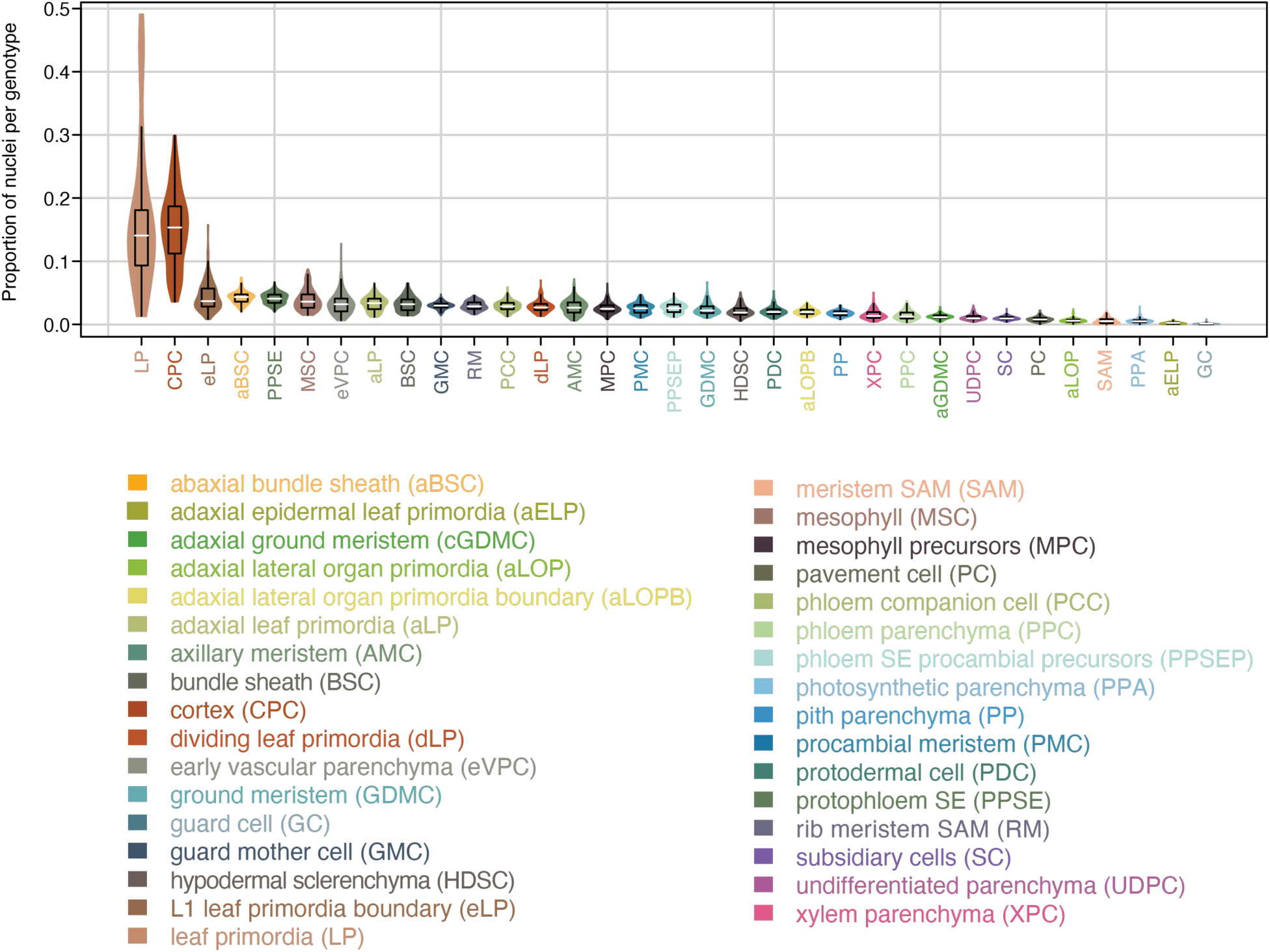
Distribution of cell type proportions among genotypes. Violin plot depicting the distribution of cell type proportions across all genotypes for each individual cell state.

**Fig. S7:**
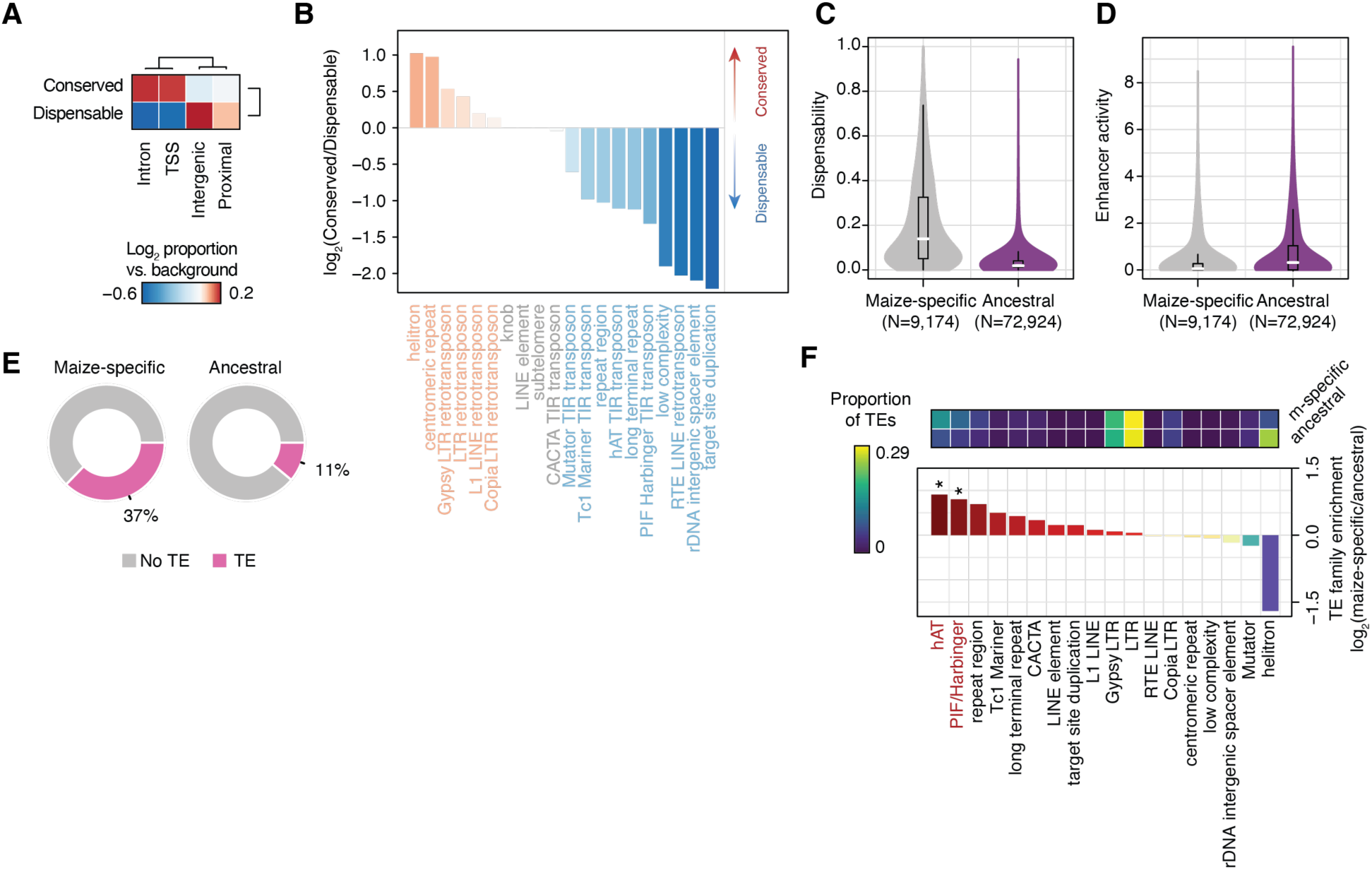
Evaluation of *cis-*regulatory variation among maize inbred lines. (**A**) Log_2_ transformed proportion of conserved and dispensable ACRs compared to all ACRs across different genomic contexts. (**B**) Log_2_ transformed ratio of conserved to dispensable ACRs proportions overlapping various classes of repetitive elements. (**C**) Distribution of dispensability (fraction of genotypes with WGS alignments covering less than 10% of the ACR) for maize-specific (grey) and ancestrally conserved (purple) ACRs. (**D**) Distribution of enhancer activity (log_2_[RNA/DNA] from STARR-seq) for maize-specific (grey) and ancestrally conserved (purple) ACRs. (**E**) Distribution of maize-specific (left) and ancestrally conserved (right) ACRs overlapping transposable elements by at least 50% of the ACR. (**F**) Top, proportion of ACRs overlapping various classes of transposable elements split by maize-specific and ancestrally conserved. Bottom, log_2_ enrichment of various transposable element families in maize-specific ACRs compared to ancestrally conserved ACRs.

**Fig. S8:**
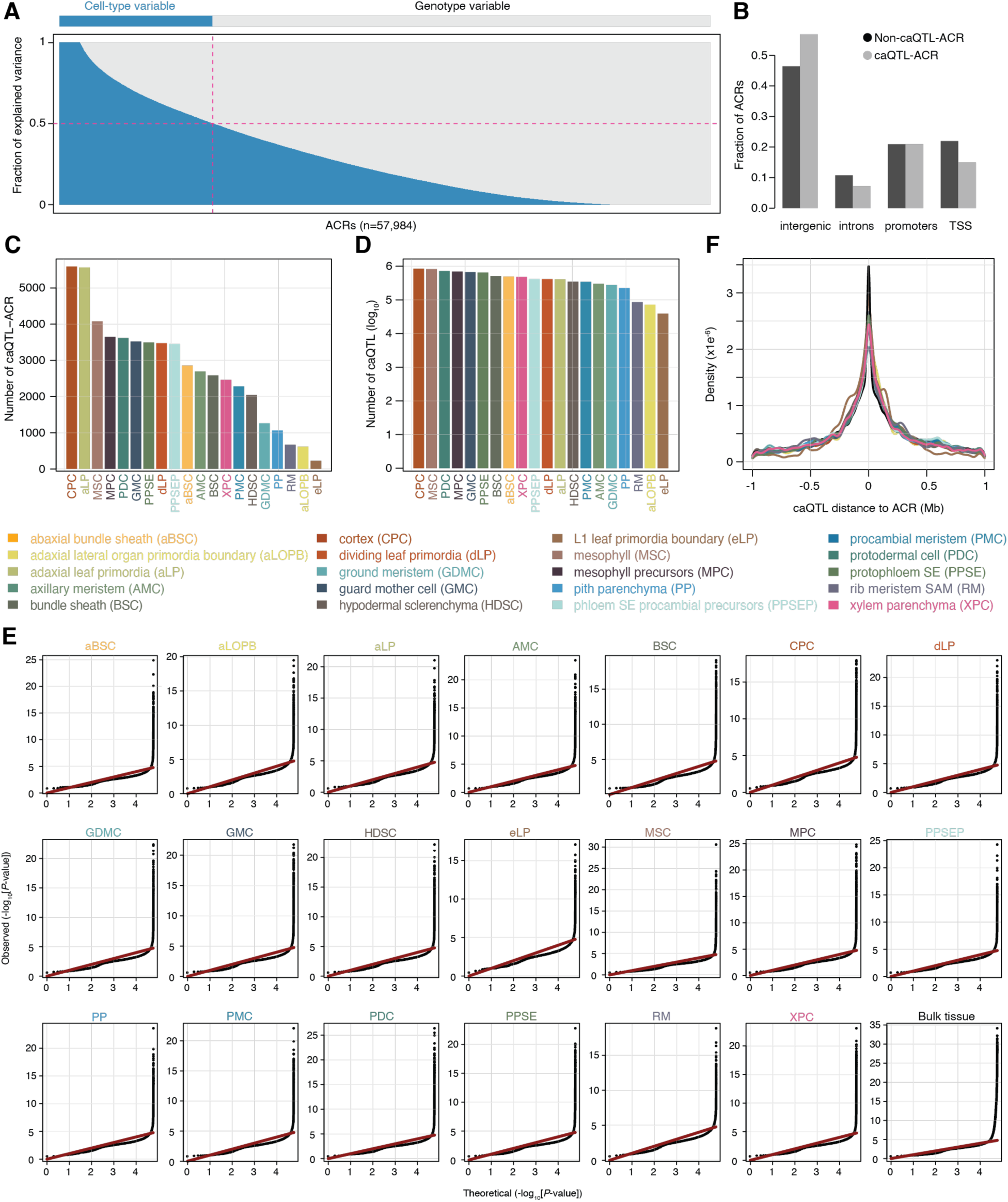
Distribution of caQTL and caQTL-ACRs across cell states. (**A**) Distribution of the relative fraction of variance explained by cell state or genotype for each ACR. (**B**) Fraction of caQTL-ACRs (grey) and non-caQTL-ACRs corresponding to various genomic contexts. (**C**) Barplot of the total counts of caQTL-ACRs by cell state. (**D**) Barplot of the log_10_ transformed counts of caQTL by cell state. (**E**) Quantile-quantile plot of nominal *P*-values of the lead SNV per ACR from caQTL mapping across 20 distinct cell states and bulk tissue. (**F**) Distribution of caQTL distances relative to the cognate ACR across all cell states.

**Fig. S9:**
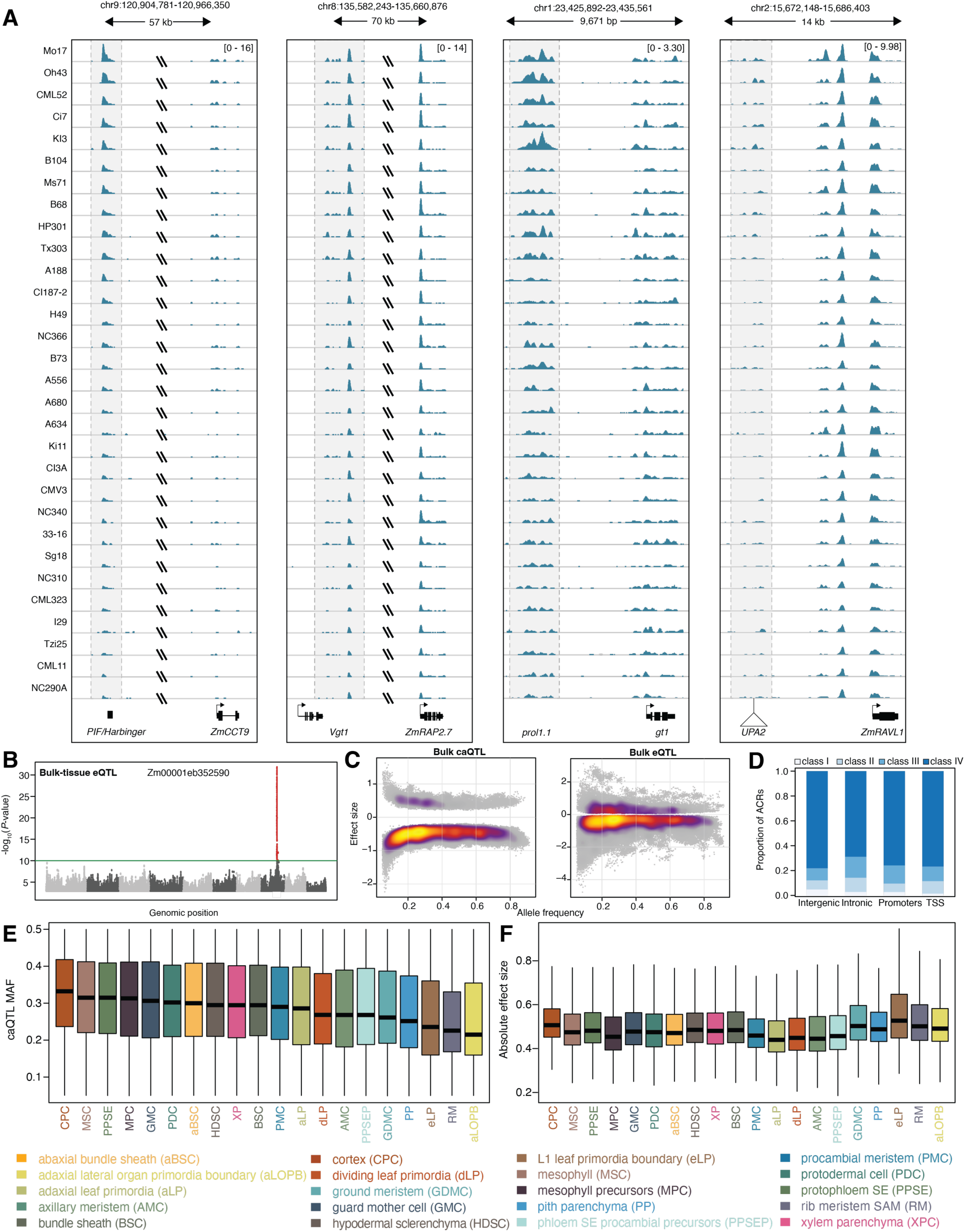
caQTL identifies known regulatory variants. (**A**) Genome browser view of four *cis*-regulatory variants with known trait effects. (**B**) Manhattan plot of eQTL analysis for an exemplary gene. (**C**) Scatter density plot illustrating the relationship between alternate allele frequency (x-axis) and effect size (y-axis) for bulk tissue caQTL (left) and bulk tissue eQTL (right). (**D**) Proportion of caQTL/eQTL effect classifications in intergenic, intronic, promoter, and TSS-localized ACRs. (**E**) Distribution of caQTL minor allele frequency by cell state. (**F**) Distribution of absolute effect sizes of caQTL grouped by cell state.

**Fig. S10:**
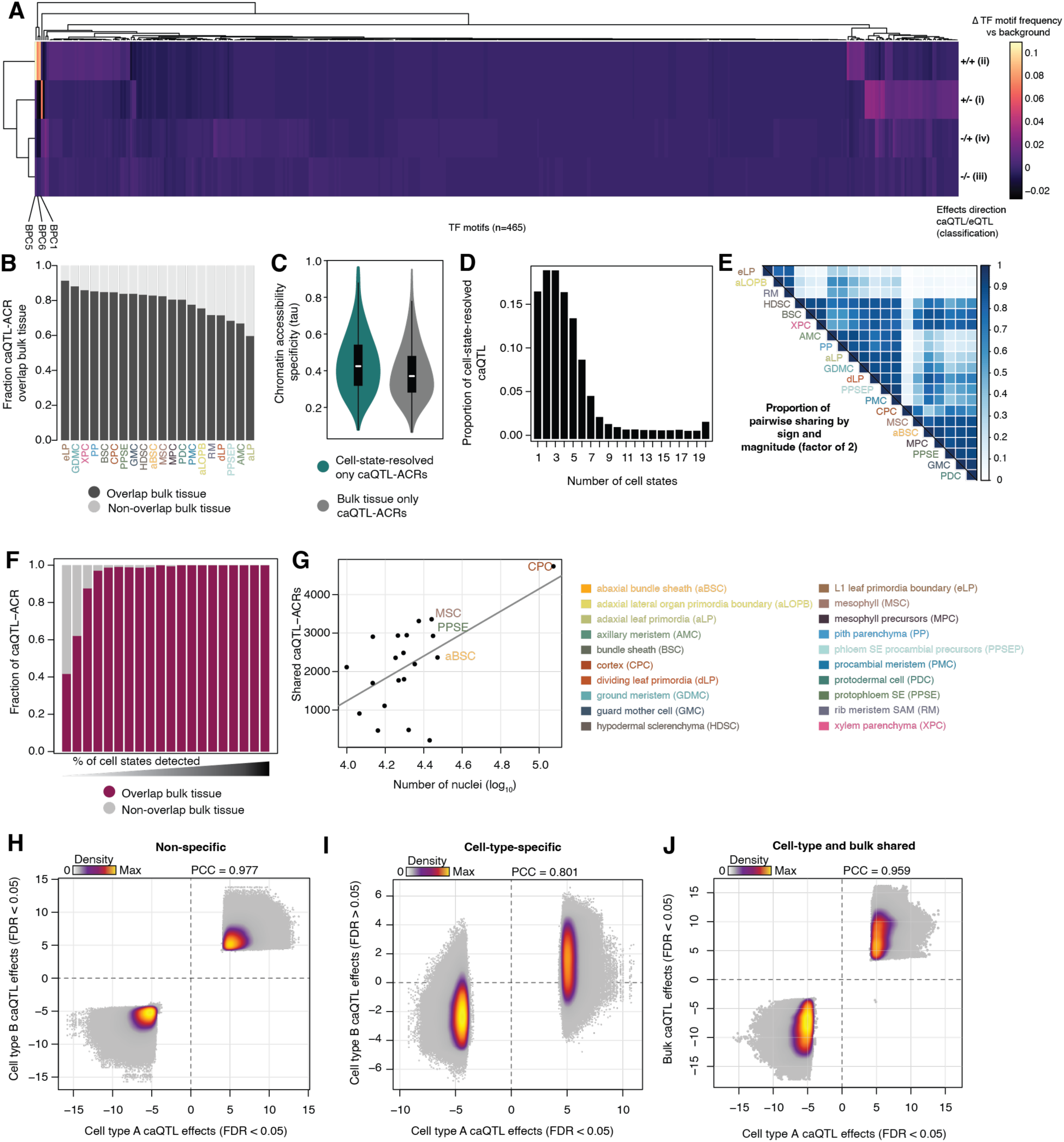
Evaluation of caQTL context specificity. (**A**) Heatmap of TFBS frequencies overlapping matched caQTL/eQTL split by effects direction classes. TFBS frequencies across all ACRs (background) was subtracted from each classification. (**B**) Fraction of cell-state-resolved caQTL-ACRs that overlap caQTL-ACRs from bulk tissue. (**C**) Distribution of ACR specificity scores (Tau) for cell-state-resolved only caQTL-ACRs and bulk tissue only caQTL-ACRs. (**D**) Distribution of caQTL sharing among cell states (within a factor of 2 and the same sign). (**E**) Pairwise fraction of shared caQTL among cell states. (**F**) Fraction of cell-state resolved caQTL-ACRs that overlap bulk tissue caQTL-ACRs split by the number of cell states in which a caQTL- ACR was detected. (**G**) Scatter plot of the number of shared caQTL-ACRs between an individual cell state and bulk tissue (y-axis) against the number of nuclei identified from a given cell state (x-axis). The top four most abundant cell states are labeled. (**H**) Density scatterplot of pairwise comparison of standardized effects for the same caQTL found in at least two cell types. (**I**) Density scatterplot of pairwise comparison of caQTL standardized effects for cell-state-specific caQTL in the focal cell state (x-axis, caQTL FDR < 0.05) compared to all other cell states (y-axis, caQTL FDR > 0.05) in a pairwise fashion. (**J**) Density scatterplot of pairwise comparison of caQTL standardized effects for overlapping cell-state-resolved and bulk tissue caQTL.

**Fig. S11:**
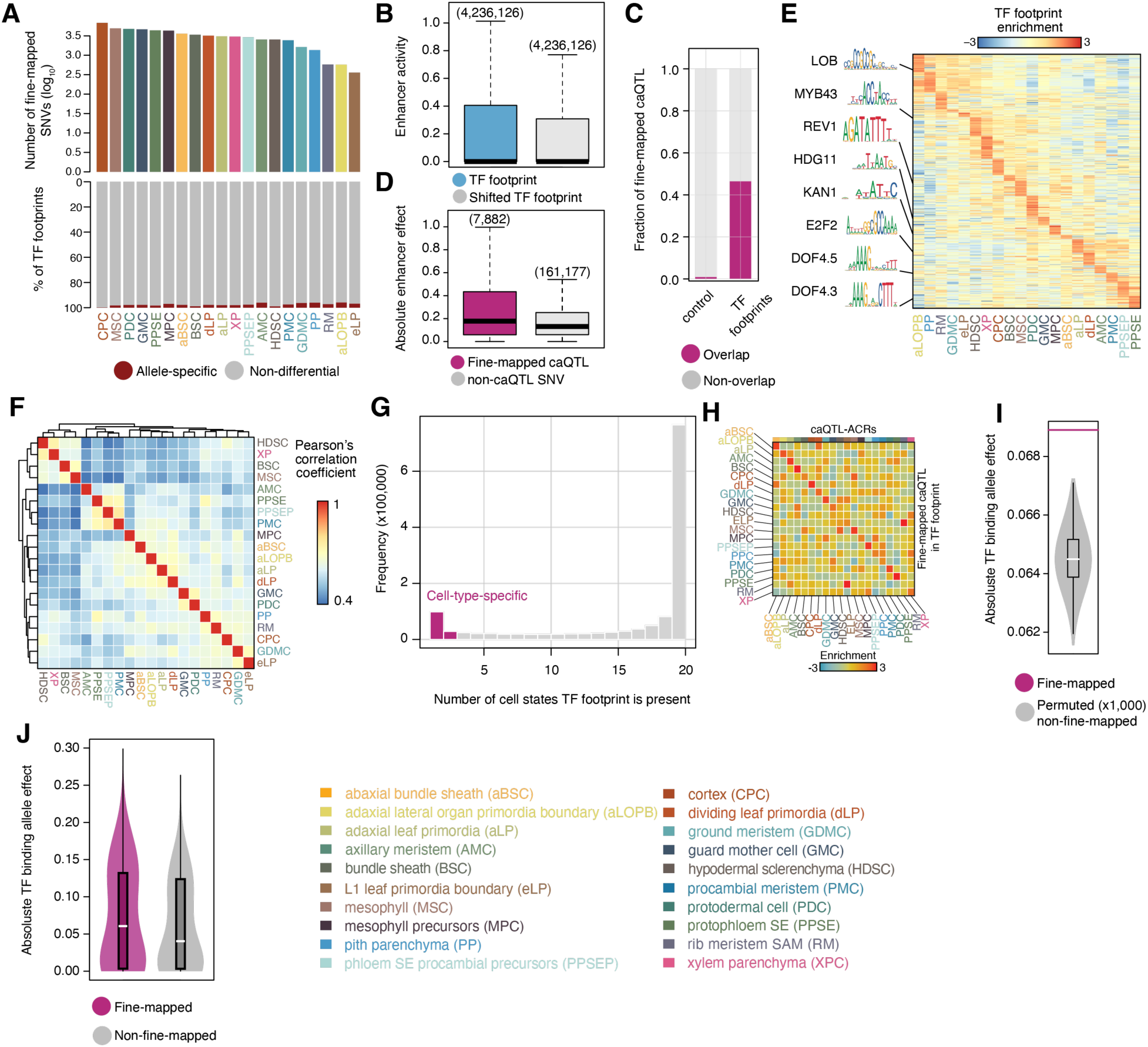
Fine-mapped caQTL effect TFBSs. (**A**) Top, barplot of the log_10_-transformed number of fine-mapped caQTL identified in each cell state. Bottom, fraction of allele-specific and non-allele-specific TF footprints across 20 cell states. (**B**) Fraction of fine-mapped caQTL overlapping TF footprints (right) or randomized control regions (left) within the same ACRs and equal number of nucleotides. (**C**) Distribution of enhancer activity overlapping TF footprints (blue) and control regions (TF footprints shifted by 50-bp). (**D**) Distribution of enhancer allelic effects for fine-mapped caQTL (left, purple), and non-caQTL variants (right, grey) located within TF footprints. (**E**) Heatmap of TF footprint enrichment for various TF motifs (rows) across 20 cell states (columns). Representative TF motifs are illustrated on the left. (**F**) Pair-wise Pearson’s correlation coefficients of TF footprint enrichment between cell states. (**G**) Distribution of TF footprint frequency across cell states. Pink bars represent cell-state-specific TF footprints (TF footprints found in two or less states). (**H**) Cell-type-specific enrichment of fine-mapped caQTL embedded TF footprints with caQTL-ACRs. (**I**) Distribution of permuted absolute TF affinity allele effects from non-caQTL SNVs overlapping TFs (grey violin plot) matched to the same number of fine-mapped variants. The average TF affinity allele effect of fine-mapped variants is shown as a pink line. (**J**) Distribution of TF affinity allele effects for fine-mapped caQTL (pink) and non-caQTL SNVs (grey).

**Fig. S12:**
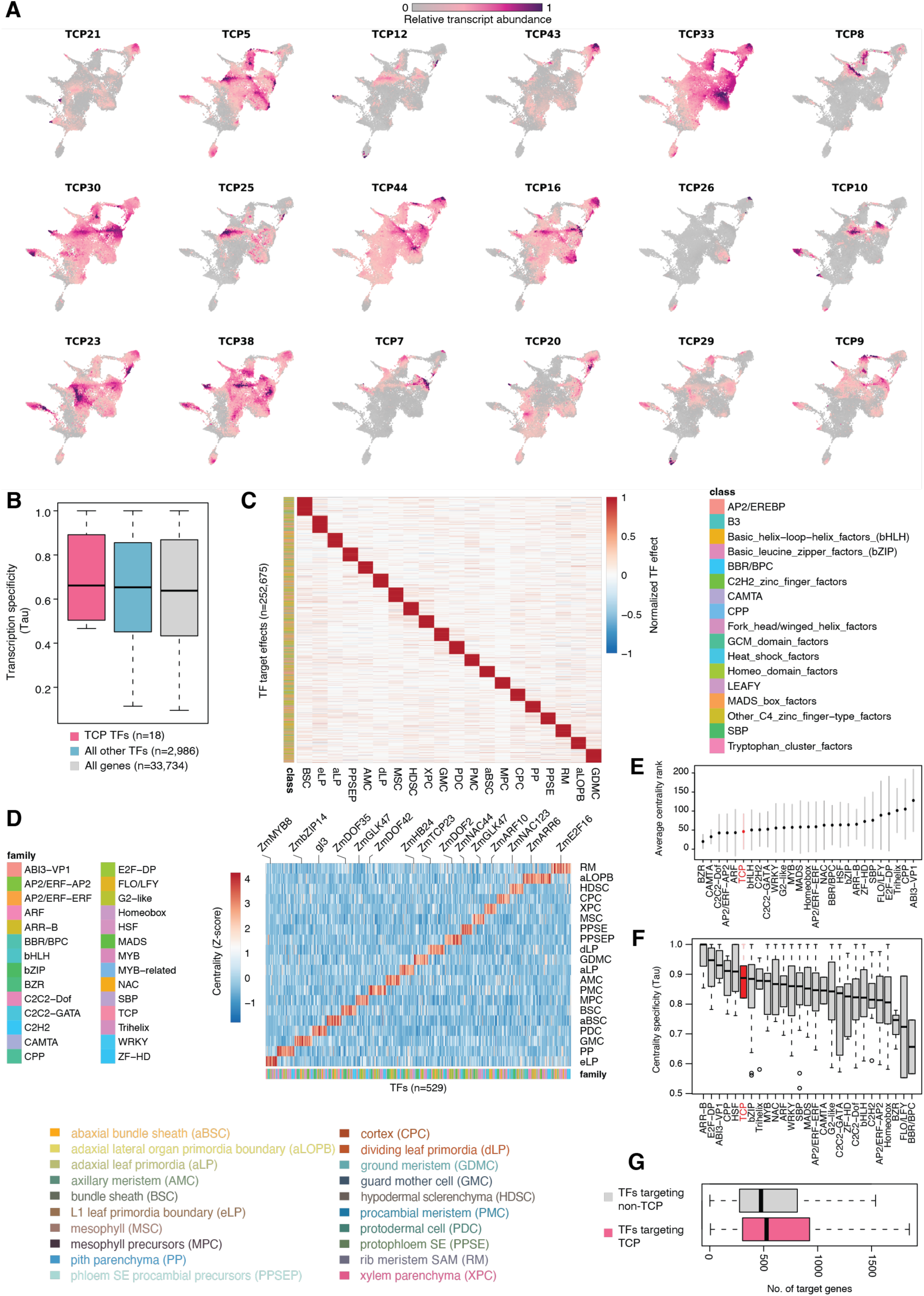
Regulatory patterns for TCP transcription factors. (**A**) UMAP nuclei embedding colored by TCP transcript abundances. (**B**) Distribution of TF transcript abundance specificity. (**C**) TF model effect estimates (scaled from -1 to 1 for each feature) per ACR across all TFs for each cell state. (**D**) Normalized (Z-score) centrality scores for each TF in each cell state. (**E**) Mean and standard deviation of the minimum centrality rankings across cell states by TF family. (**F**) Distributions of centrality score specificity (tau) by TF family. (**G**) Distribution of target counts for TFs associated with ACRs upstream of TCPs (pink) and all other genes (grey).

**Fig. S13:**
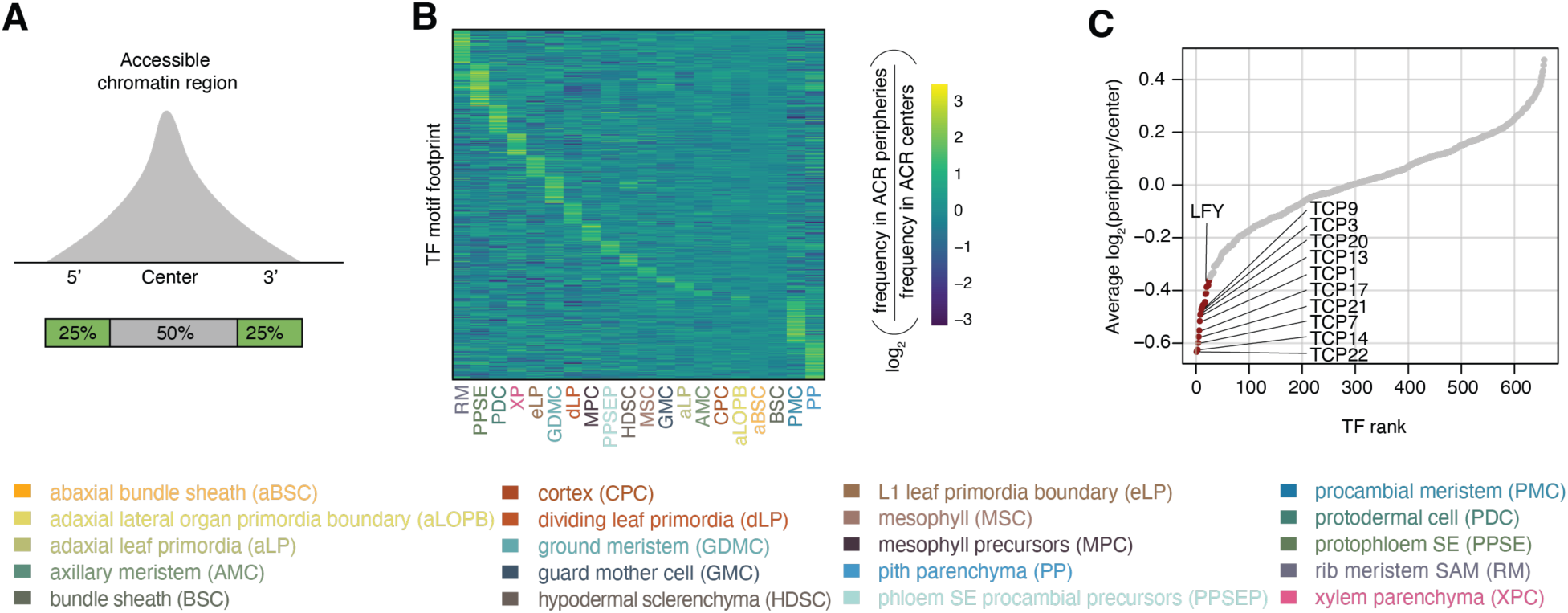
Unique TFBSs are enriched in dynamic ACR boundaries: (**A**) Illustration of TF footprint assignments relative to ACR coordinates. (**B**) Heatmap of the log_2_ enrichment of TF footprints (rows) in ACR peripheries or ACR centers across cell states (columns). (**C**) Ranked scatter plot of the average log_2_ enrichment of TF footprints near ACR centers.

**Fig. S14:**
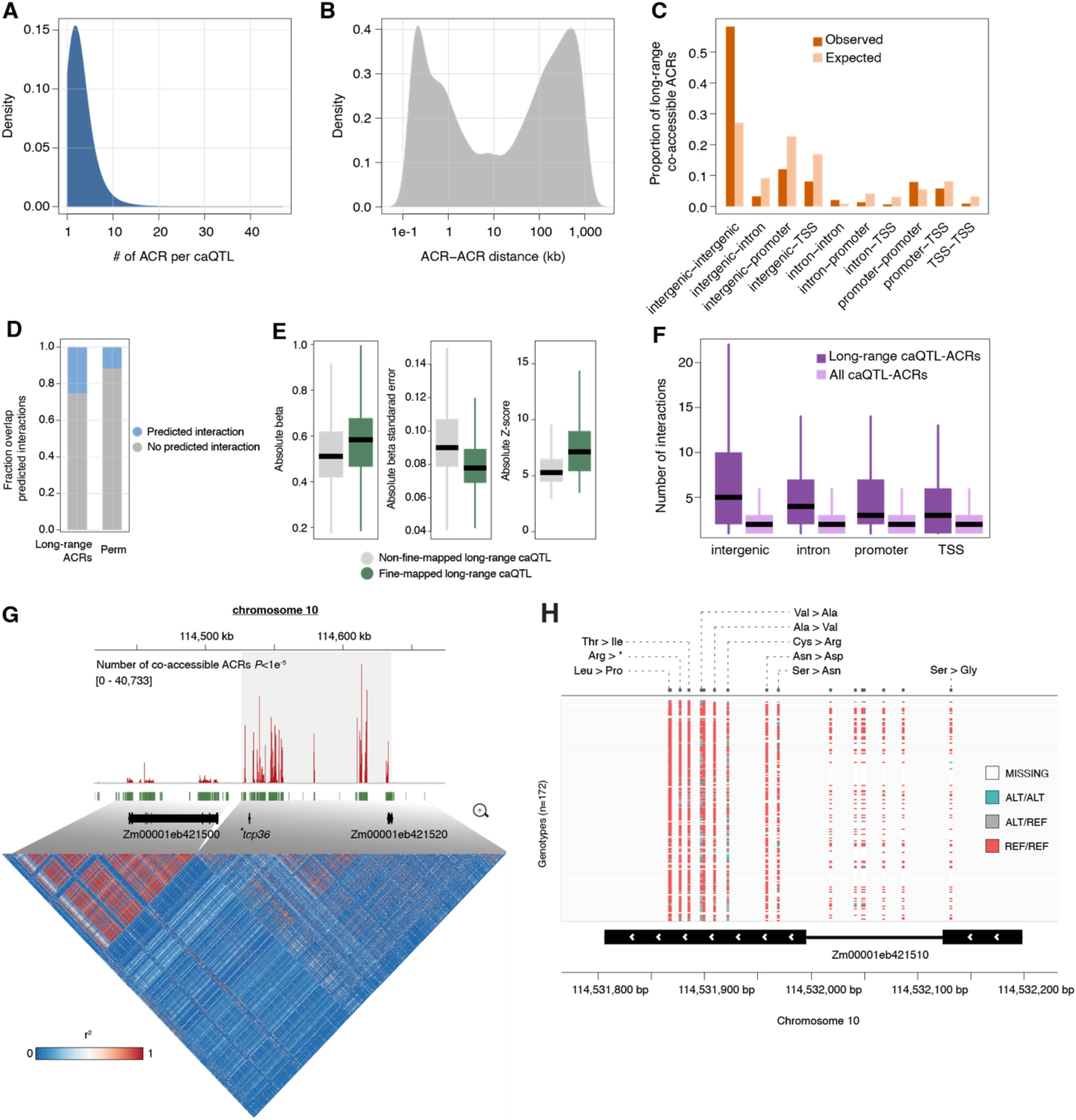
Evaluation of long-range caQTL. (**A**) Distribution of ACRs associated with the same caQTL variant. (**B**) Distribution of ACR-ACR distances (kb) for long-range ACRs associated with the same caQTL. (**C**) Proportion of long-range co-accessible ACRs corresponding to various genomic contexts split by observed long-range caQTL-ACRs (observed) and all possible ACR-ACR interactions (expected). (**D**) Proportion of long-range ACRs and permuted interactions overlapping predicted interactions from the deep learning model, GenomicLinks. (**E**) Left, distribution of absolute effect sizes; middle, distribution of absolute effect standard errors; right, distribution of absolute Z-scores. Grey, non-fine-mapped long-range caQTL. Green, fine-mapped long-range caQTL. (**F**) Distribution of long-range interaction counts by genomic context for long-range caQTL-ACR (the ACR containing the local caQTL; observed) and a random selection of the same number of ACR-ACR links derived from all caQTL-ACRs (expected). (**G**) Top: Genome-browser view of co-accessible ACR hits per SNV (red bars) in a 250-kb window surrounding *tcp36.* Green squares indicate the positions of individual SNVs. Bottom: Heatmap of linkage-disequilibrium (r^2^) among all SNVs within the ∼250-kb window. (**H**) Genome browser view of *tcp36* missense variants and genotype frequency across the 172 maize inbreds.

**Fig. S15:**
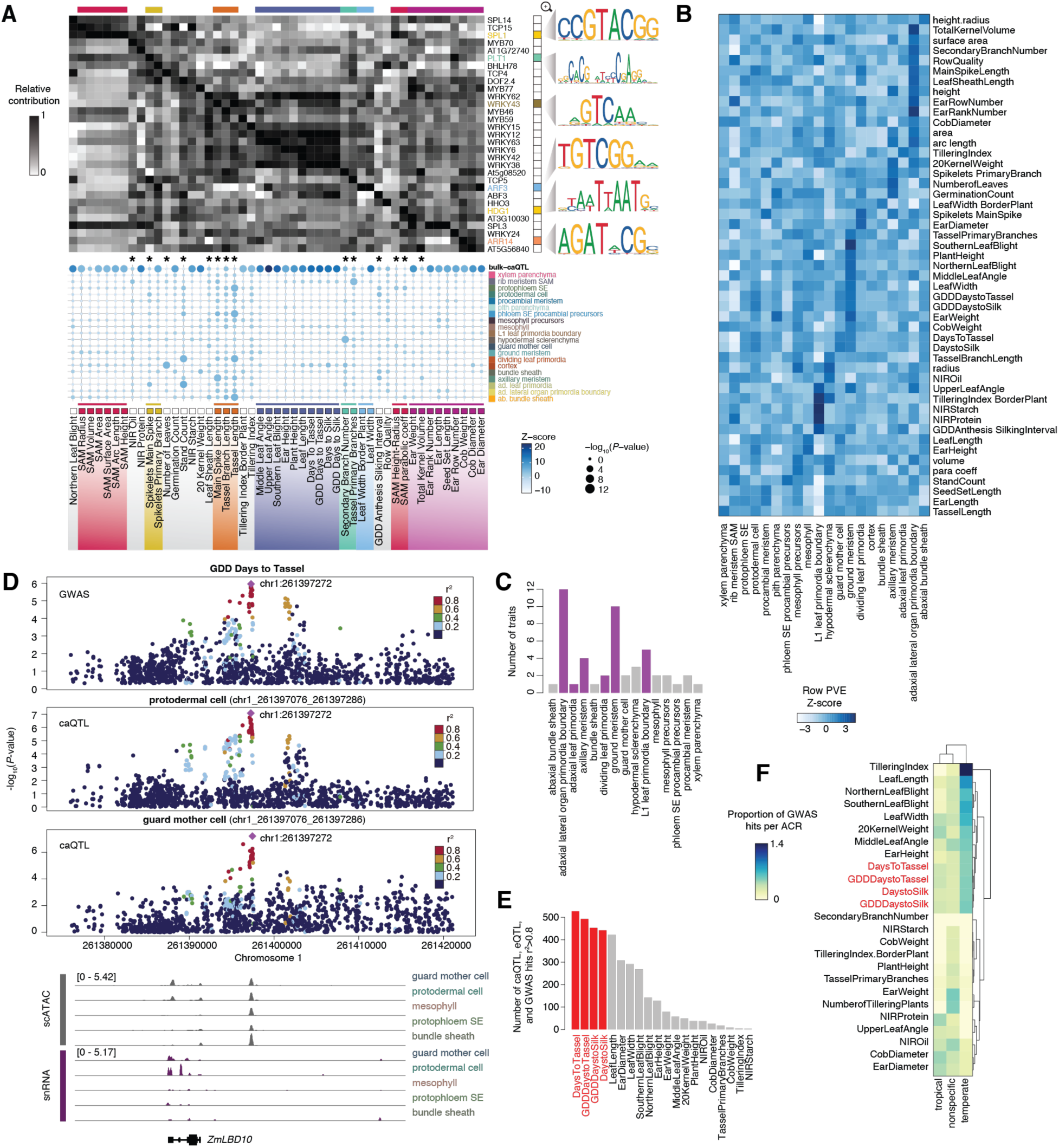
caQTL explain phenotypic variance. (**A**) Top, heatmap of the top aggregated TFBS effects by phenotype. Select motifs are shown on the right. Bottom, bubble plot of percent variance explained (permutation test; 10,000x) by bulk-caQTL (top row) and caQTL identified only by cell-state-resolving profiling (all other rows). Asterisks indicate traits where cell-state-resolved caQTL explain significantly more trait variation than bulk-caQTL. (**B**) Heatmap of standardized mean per caQTL percent of variance explained for 49 traits across 20 cell states. (**C**) Bar plot of the number of traits where the cell state had the largest proportion of variance explained. (**D**) Top; manhattan plots for GDD Days to Tassel and chromatin accessibility in two epidermal cell states. Bottom; cell state-aggregated ATAC-seq and RNA-seq coverages for four cell states. (**E**) Counts of GWAS hits in LD with shared caQTL and eQTL. (**F**) Fraction of all GWAS hits mapping to temperate, tropical, and non-subpopulation specific ACRs, scaled by the number of ACRs.

**Fig. S16:**
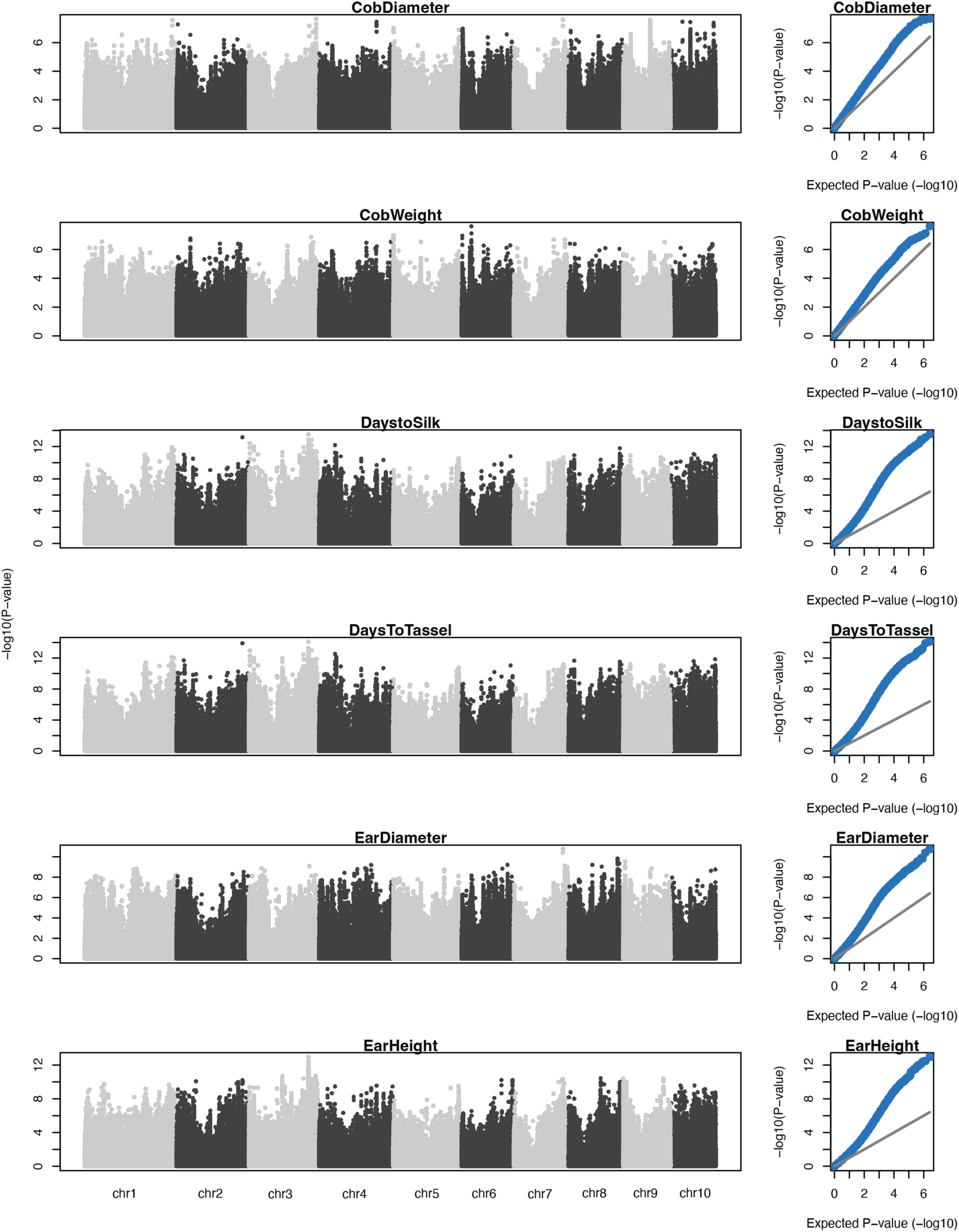
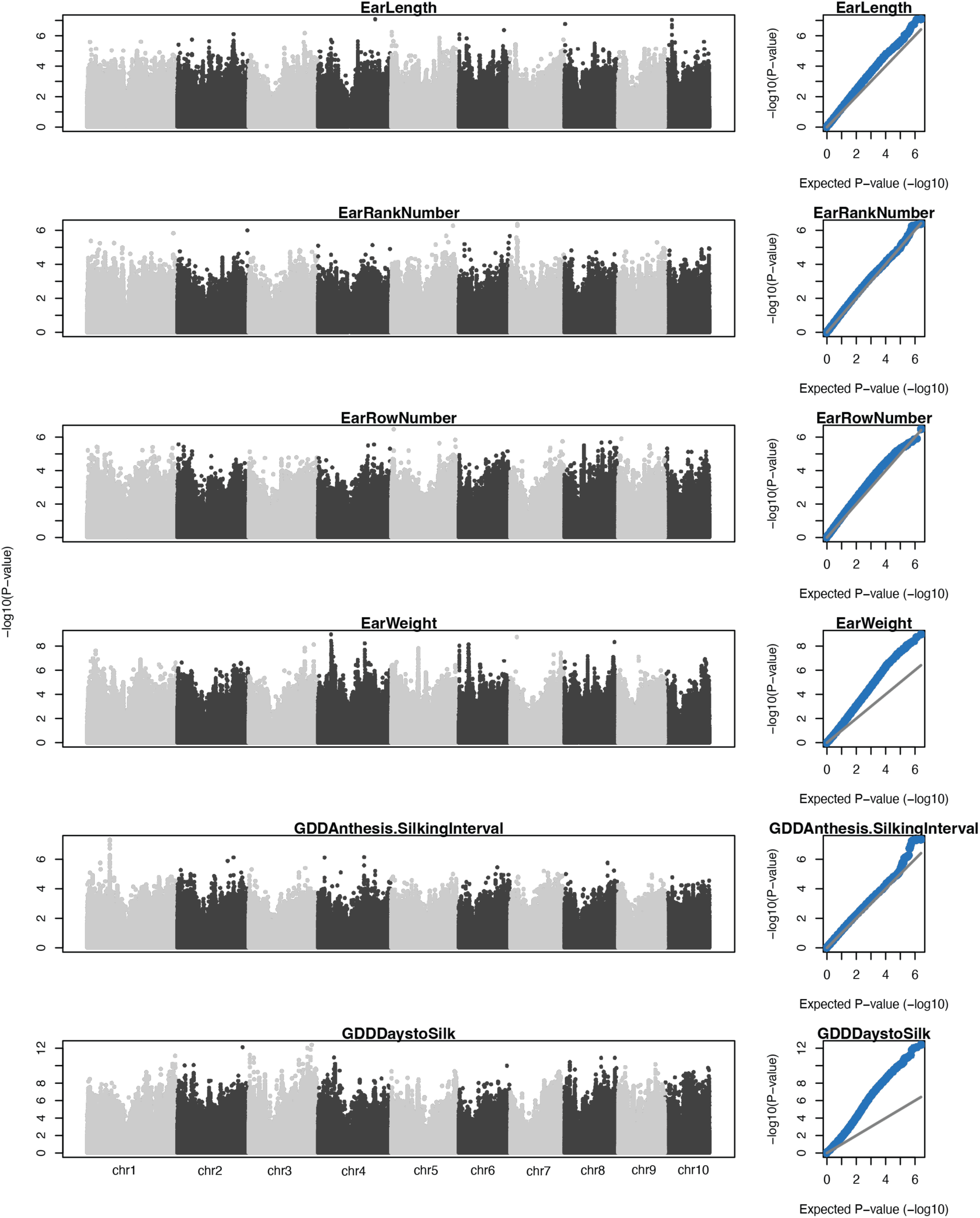
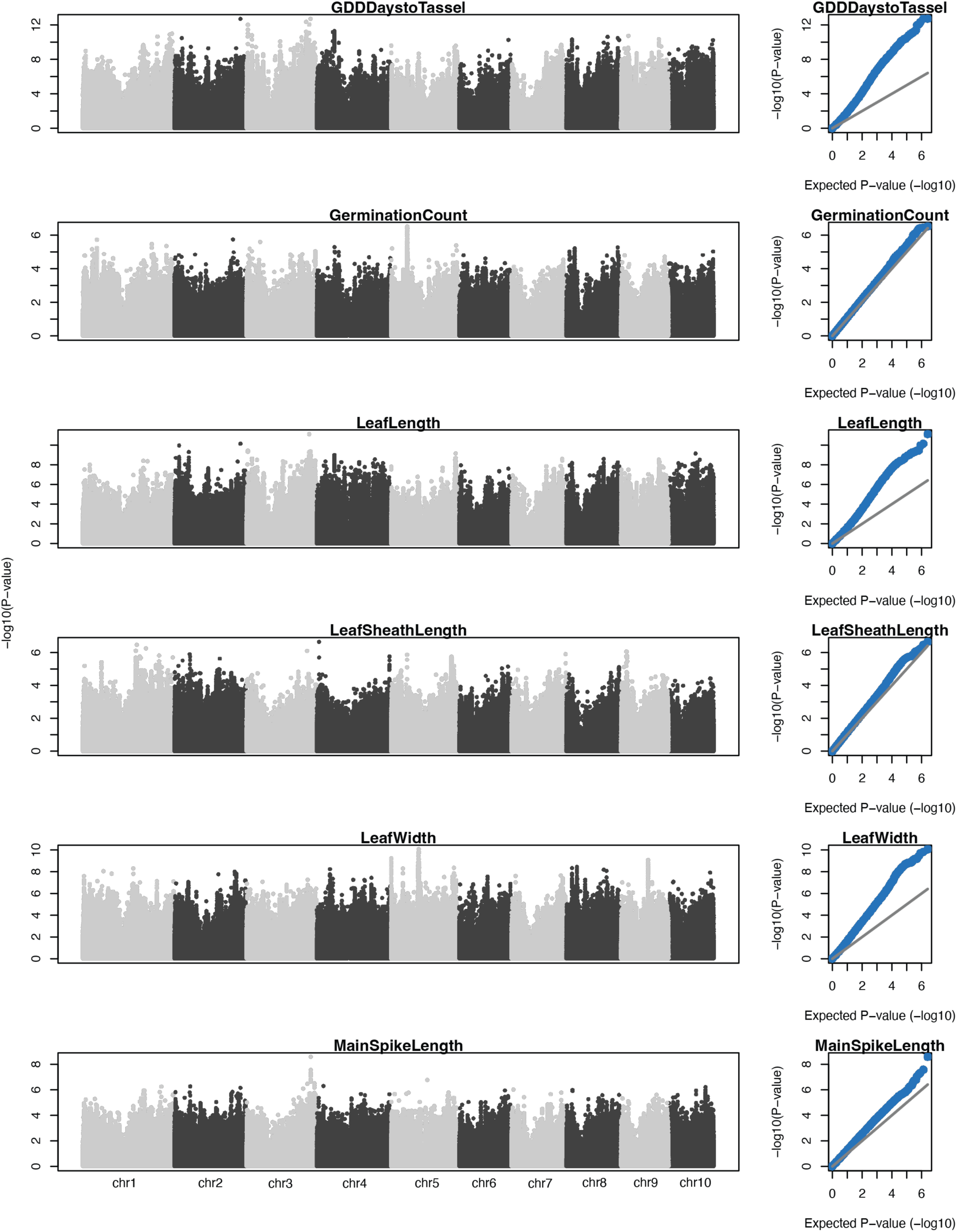
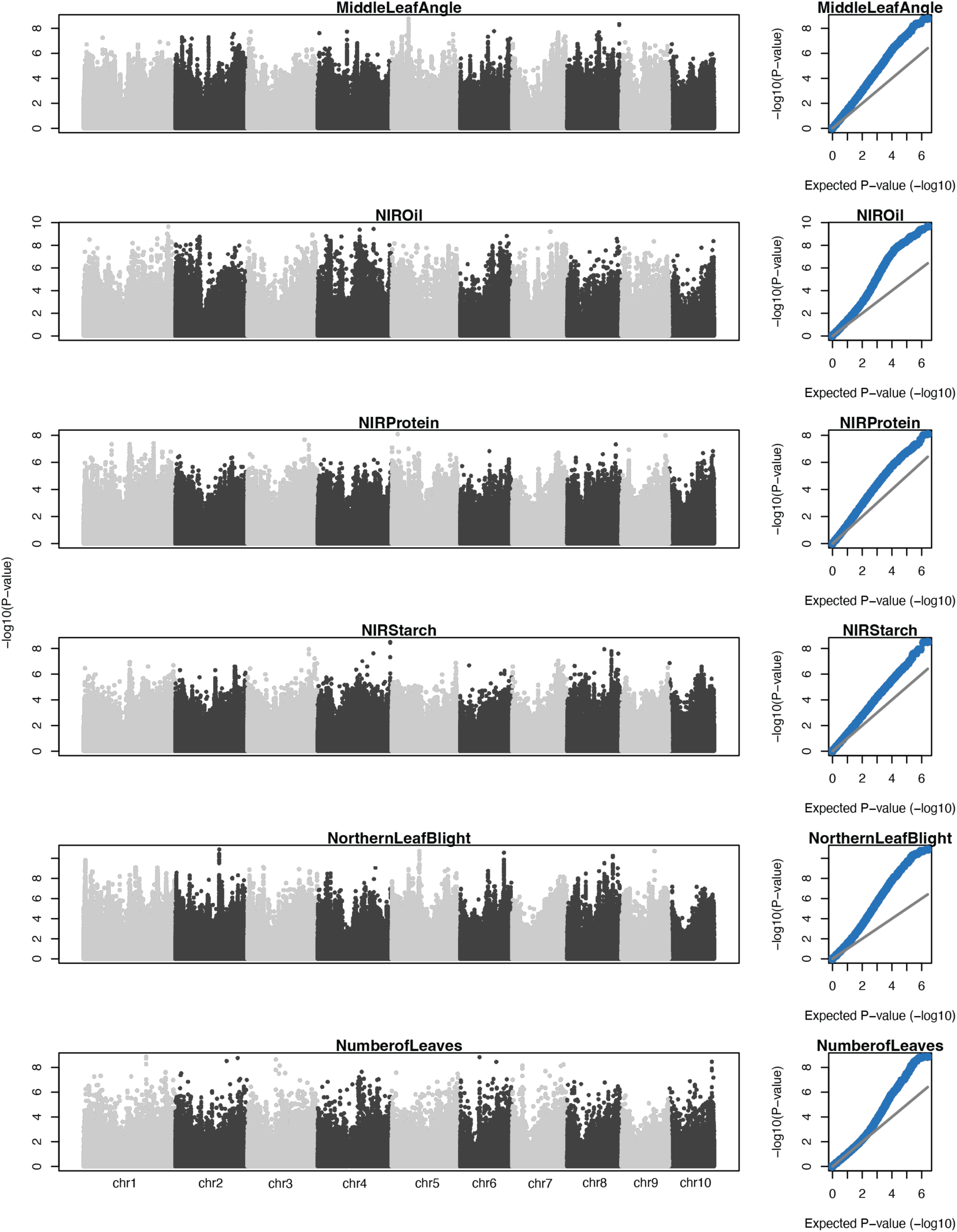
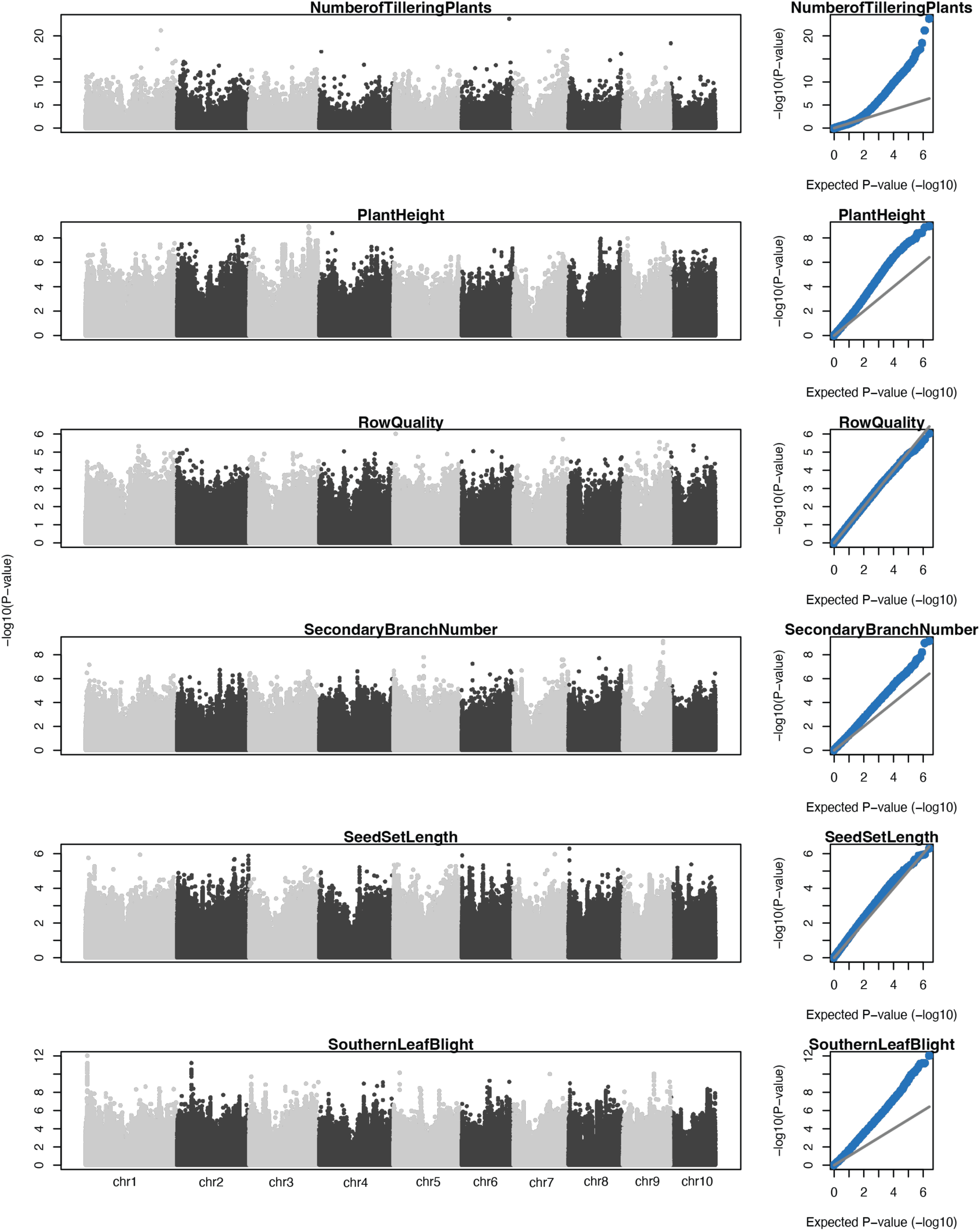
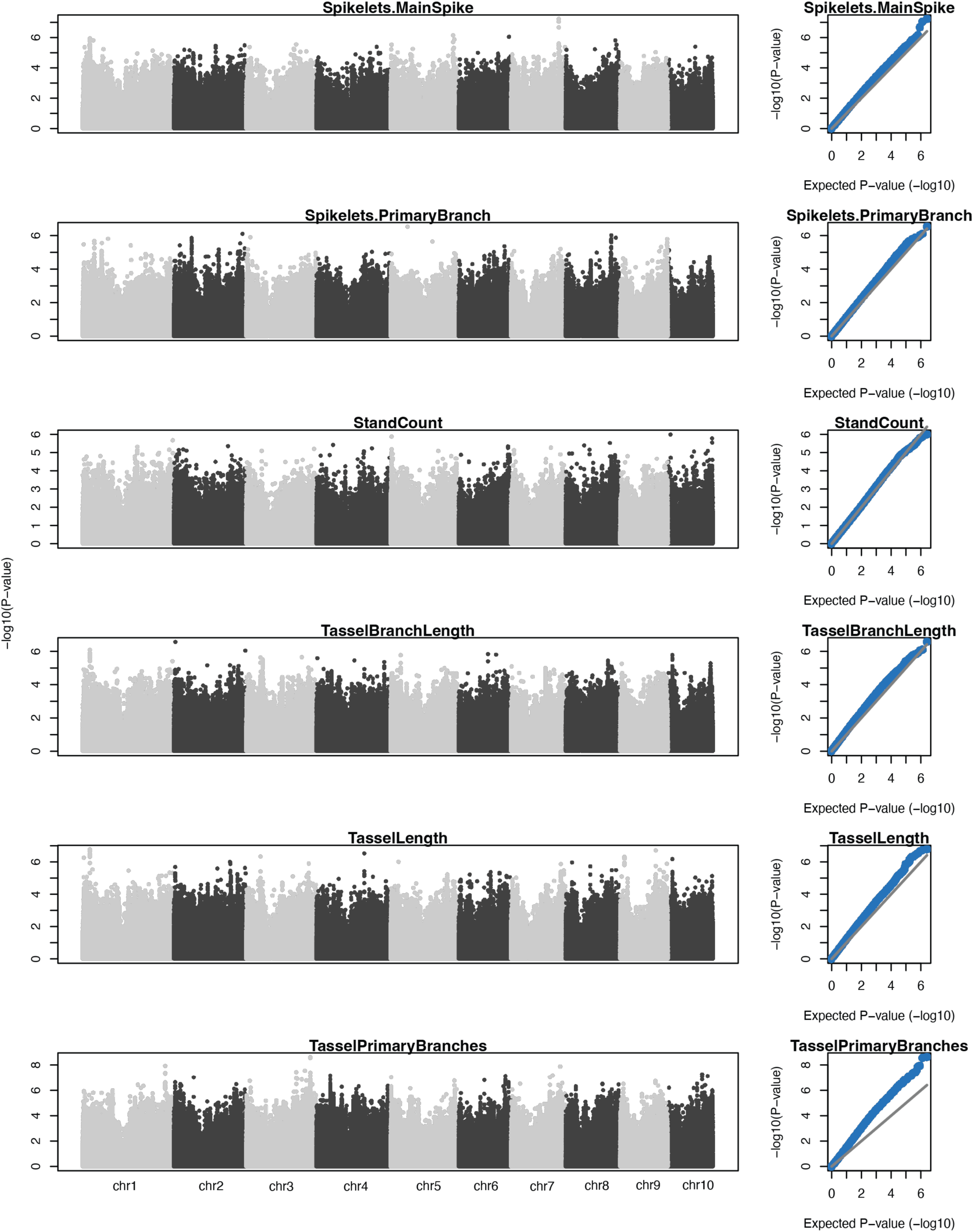
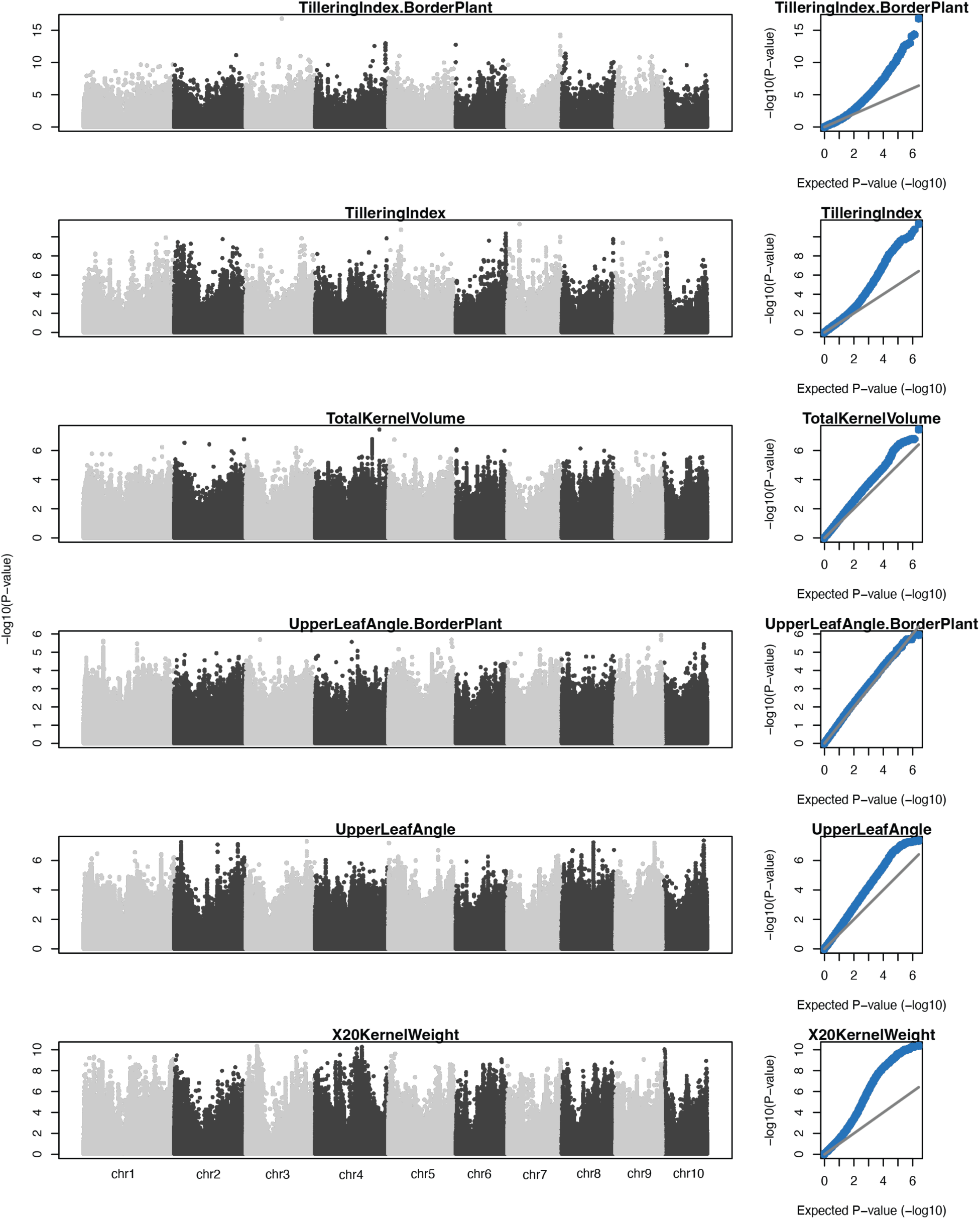
Manhattan and Q-Q plots from GWAS of the Goodman-Buckler Association Panel (*n*=277)

## Captions for additional separate files

Data S1. (separate file)

scATAC-seq library summary statistics.

Data S2. (separate file)

Genotype pooling strategy.

Data S3. (separate file)

Accessible chromatin regions classifications.

Data S4. (separate file)

Motif conservation summary statistics.

Data S5. (separate file)

Fine-mapped caQTL summary statistics.

Data S6. (separate file)

Co-accessibility *trans-*QTL mapping candidate gene list.

Data S7. (separate file)

Integration of caQTL, eQTL, and GWAS hits.

Data S8. (separate file)

Differentially accessible subpopulation ACRs.

Data S9. (separate file)

Motif enrichment in subpopulation-specific ACRs.

Data S10. (separate file)

Motif enrichment overlapping high and low *F*_st_ caQTL.

